# Expression of *AtCAN1* and *AtCAN2* genes of the plant SNc nuclease family correlates with programmed cell death and endoreduplication, indicating their role in the recycling of nucleic acid components

**DOI:** 10.1101/2025.10.10.680537

**Authors:** Rafal Krela, Elzbieta Poreba, Krzysztof Lesniewicz

**Affiliations:** Department of Plant Physiology, Faculty of Biology, Adam Mickiewicz University, Uniwersytetu Poznańskiego 6, Poznań, 61-614, Poland; Department of Genetics, Faculty of Biology, Adam Mickiewicz University, Uniwersytetu Poznańskiego 6, Poznań, 61-614, Poland; Department of Molecular and Cellular Biology, Faculty of Biology, Adam Mickiewicz University, Uniwersytetu Poznańskiego 6, Poznań, 61-614, Poland

**Keywords:** Staphylococcal-like nuclease, DNA degradation, programmed cell death, polyploidy, endoreduplication, pathogen infection, GUS assay, *Arabidopsis thaliana*

## Abstract

**Background:** Controlled degradation of genomic DNA is a common feature of all known cases of programmed cell death (PCD) in animals and plants. In plants, nucleic acid degradation during PCD facilitates the redistribution of their constituent building blocks. Previous studies have shown that nucleases from the S1/P1 family are involved in this process; however, due to the complexity of the process, it has been hypothesized that nucleases from other families, including staphylococcal-like nucleases (SNc), may also participate. In *Arabidopsis*, this family comprises two nucleases with atypical plasma membrane localization for this enzyme class: AtCAN1 and AtCAN2.

**Results:** Using the promoter-driven GUS reporter assay, we showed that genes encoding SNc nucleases are expressed in tissues grouped into three main categories. The first category includes plant structures that are clear examples of organs undergoing PCD, such as the root cap, vascular bundle elements, the tapetum, maturing seed pods, and senescent leaves. The second category comprises cells whose function involves interaction with the external environment and which are susceptible to pathogen attack. This group includes root hairs, stomatal guard cells, and hydathodes. The third group of plant structures showing SNc nuclease activity consists of elements characterized by endoreduplication, i.e., stipules, trichomes, and the basal parts of the hypocotyl. The analysis of microarray and RNA-seq transcriptomic data further confirmed the expression of both SNc genes in these three categories. Moreover, we presented the effects of a mutation that eliminates AtCAN1 expression on plant development.

**Conclusions:** Our studies show that SNc nucleases are as broadly involved in the DNA degradation during plant PCD as the previously reported S1/P1 proteins. The frequent overlap in their expression profiles suggests cooperative action. Whereas SNc nucleases localize to the plasma membrane, S1/P1 nucleases are nuclear, indicating distinct yet complementary nucleolytic pathways. We further demonstrate that SNc nucleases are specifically expressed in organs that do not undergo PCD but are characterized by endoreduplication, implicating them in an unexplored mechanism for redistributing polyploid DNA components.

## Background

The development of multicellular eukaryotic organisms is inextricably linked to degradative processes that occur both in living cells, such as autophagy, and in cells destined for elimination through various forms of programmed cell death (PCD). At the molecular level, these processes primarily target membrane structures, proteins, and nucleic acids. In plants, a wide range of PCD processes has been documented. Model examples include developmental events such as xylem differentiation, root cap growth, tapetum degradation associated with pollen development, and senescence, which has been studied most extensively in withering leaves [1]. Degradative processes are also observed in environmentally induced PCD. The best-characterized case is the hypersensitive response (HR), a classical defence mechanism in which plant cells undergo localized death upon pathogen attack [2].

In all studied cases of PCD, intense nucleolytic activity has been observed, leading to the degradation of both genomic DNA and cellular RNA. Although DNA elimination occurs during PCD in both plants and animals, the underlying biological function differs substantially. In animals, DNA degradation primarily serves to protect surrounding tissues from inflammation. Such inflammation could otherwise be triggered by the release of nucleic acids from disintegrating cells into the intercellular space [3]. In contrast, this explanation does not apply to plants. Instead, it is generally assumed that nucleic acid degradation in plant tissues undergoing PCD facilitates the redistribution of their constituent building blocks. This role is particularly important because nucleic acids are rich in reduced nitrogen and readily available phosphorus - two essential mineral nutrients that frequently limit plant growth due to their scarcity in the natural environment. Consequently, the efficient recycling and redistribution of these elements is considered crucial for sustaining plant development [4].

The problem of DNA component redistribution is also connected with the phenomenon of DNA endoreduplication. Endoreduplication occurs in various plant tissues, but its biological significance is still unclear. However, considering how many valuable building blocks polyploid DNA contains, the question arises about their final fate, i.e., whether they can be recycled. In storage tissues such as endosperm and cotyledons, where endoreduplication occurs [5] and where, during the mobilization of accumulated reserves, strong nucleolytic activity has also been reported [6, 7], it has been proposed that the accumulated DNA may serve as a source of elements for the synthesis of nucleic acids in developing organs [8]. However, it remains unclear whether polyploid DNA in tissues that do not serve storage functions can also be utilized for a similar purpose.

A group of hydrolytic enzymes, commonly referred to as degradative nucleases, is responsible for the nonspecific breakdown of nucleic acids. Unlike nucleases involved in DNA repair or recombination, degradative nucleases exhibit high catalytic activity, low substrate specificity, and tissue-specific expression. Plants and animals use enzymes from distinct protein families to perform this function. In plants, BFN1, a member of the S1/P1 endonuclease family, is the nuclease for which the most compelling evidence supports its role in PCD [9].To establish this function, studies employing the GUS reporter gene under the control of the BFN1 promoter proved crucial [10]. In addition, analysis of BFN1 gene expression profiles from available microarray data confirmed that this nuclease’s expression correlates with known instances of PCD [11].

However, analysis of degradative nucleolytic activities detected in various plant tissues undergoing PCD suggests that multiple nucleases are likely involved in these processes [6]. These enzymes differ considerably in their molecular weight and catalytic requirements. As we have established, this diversity arises primarily from the fact that individual proteins of the S1/P1 family are catalytically active under different pH conditions - mainly acidic or neutral - and also from the fact that they utilize distinct divalent cations, most commonly Zn^2+^, Ca^2+^, or Mn^2+^, as cofactors. Moreover, members of this family are further distinguished by their sensitivity to inhibition by various factors [11]. Such extensive diversity among degradative nucleases can be attributed to their roles in different forms of PCD that occur in tissues exhibiting substantial morphological and physiological variation. Furthermore, because DNA degradation appears to be a multistep process, even a single type of PCD may involve the coordinated action of several nucleases, each operating under conditions specific to distinct stages. Taken together, these observations indicate that plants require a broad repertoire of nucleases capable of hydrolyzing nucleic acids under diverse conditions.

According to our hypothesis, another family of DNases that may, alongside the S1/P1 nucleases, participate in PCD-dependent DNA degradation in plants is the Staphylococcal-like nuclease family (SNc). Our previous studies on two plant members of this family, encoded by *Arabidopsis thaliana* – AtCAN1 and AtCAN2 – have demonstrated that these enzymes, similar to their bacterial homologs, are catalytically active at neutral pH in the presence of Ca^2+^ ions and display pronounced DNA-degrading activity. Moreover, expression analysis based on available microarray data suggested that tissue-specific expression of at least one of these genes, *AtCAN1*, may correlate with the progression of PCD. Notably, AtCAN1 exhibits several unusual and intriguing features for degradative nucleases, including membrane localization and an ABC transporter motif, suggesting its potential involvement in intercellular transport [12]. Additional support for the involvement of this protein family in PCD comes from the expression profile of the *CrCAN* gene from *Citrus reticulata*, which encodes a nuclease homologous to AtCAN1 and participates in DNA degradation during PCD of secretory cavity cells [13, 14].

Therefore, we conducted studies to characterize the tissue-specific expression profiles of SNc family nucleases, employing the GUS reporter gene under the control of the *AtCAN1* and *AtCAN2* promoters to assess whether their activity overlaps with known degradative processes occurring at distinct stages of plant development, using *Arabidopsis thaliana* plants transformed with the aforementioned transgenes. An extensive analysis of transcriptomic datasets from numerous microarray and RNA-seq experiments further supported these results. Moreover, we demonstrated how the loss of *AtCAN1* gene expression influences the phenotype of *A. thaliana*.

## Methods

### Generation of AtCAN1/AtCAN2 Promoter::GUS gene fusion constructs and preparation of transgenic plants

To generate the *AtCAN1* and *AtCAN2* promoter::GUS gene fusions, the binary vector pCAMBIA3301 (CAMBIA, Black Mountain, Australia) was modified by removing the CaMV 35S sequence. This was achieved by excision of the promoter using the restriction enzymes PstI and NcoI, followed by end-blunting and plasmid recircularization, resulting in a construct retaining only the multiple cloning site and the GUS gene.

To obtain the promoter sequences of the *AtCAN1* and *AtCAN2* genes from *A*.*thaliana* genomic DNA, nested PCR was performed. In the first stage, primers anchored in genomic regions flanking the target sequences were used. A 2.9 kb DNA fragment encompassing the *AtCAN1* promoter [At3g56170] was PCR-amplified with forward primer 5’-CACAGCAAGAGCAAGAGCAG-3’ and reverse primer 5’-AGACGCCGTGAGAATTCAAG-3’. Similarly, a 1.8 kb DNA fragment containing the *AtCAN2* promoter [At2g40410] was amplified with forward primer 5⍰-GGTGAGCAAGCAAAGAGGTC-3⍰ and reverse primer 5’-GCACAAATGAAAGCAGCAGA-3’. In the second stage, the promoter sequences were amplified from purified first-round PCR products using primers carrying restriction enzyme recognition sites to facilitate cloning into the modified pCAMBIA3301 plasmid. The *AtCAN1* promoter was amplified with *EcoR*I-forward primer 5’-AAAgaattcCTGAATTGGCAATATGATAAG-3’ and *Xba*I-reverse primer 5’-CCCtctagaTTCTTAGATTTGATTTTCAAC-3’ primers, while the At*CAN2* promoter was amplified with *Sac*I-forward primer 5’-AATgagctcGTATGGTGATGCGCGGTGC-3’ and *Xba*I*-*reverse primer 5’-CCCtctagaCTTTCACCCAATTTCAGGGAATC-3’.

Following purification, the PCR products were digested with the corresponding restriction enzymes and ligated into the modified pCAMBIA3301 vector digested with the same enzymes. The ligation mixtures were used to transform E. coli DH5α. After Sanger sequencing confirmed positive colonies, the resulting plasmids were introduced into *Agrobacterium tumefaciens* GV3101 by the freeze-thaw method for subsequent stable transformation of *A. thaliana*.

Stable transformation of *A. thaliana* plants with the At*CAN1*/At*CAN2* promoter::GUS constructs was performed using *Agrobacterium tumefaciens*-mediated transformation according to the floral dip method [15]. Transformants were selected and analyzed to establish stable transgenic lines containing the integrated GUS gene under the control of the respective nuclease promoter sequences. Homozygous lines were established, and T3 lines were used for the experiments.

### Plant growth and selection

For in vitro growth and selection of transformed plants, half-strength Murashige and Skoog (MS) solid medium was used. The pH of the medium was adjusted to 5.6-5.7 using KOH before adding Phytoagar, and the medium was sterilized by autoclaving. After that, BASTA (glufosinate-ammonium) was added at a final concentration of 15µg/ml as a selection agent. Sterilized seeds were sown on plates and PhytoCon in vitro plant culture vessels, subjected to vernalization for 48 h at 4°C in darkness, then transferred to a phytotron and grown at 22°C under a 16 h/8 h light/dark cycle, with a light intensity 120µmol/m^2^/s, and 70% relative humidity. Ten days after germination, *Arabidopsis* seedlings from selection plates were transferred to peat pellets (Jiffy Products) and, together with plants in PhytoCon, continued growing under the same conditions.

### GUS histochemical staining

Reporter gene expression was visualized by *in situ* histochemical staining. Whole seedlings, fragments, or organs (roots, stems, leaves, flowers, and siliques) were collected, placed in chilled 90% acetone, and kept until all samples were collected. Samples were then transferred to a 12-well plate (3.8 cm^2^ surface area per well) filled with staining buffer (50 mM phosphate buffer, pH 7.2, 0.2% (w/v) Triton X-100, 0.25 mM potassium ferricyanide (K_3_[Fe(CN)_6_]), 0.25 mM potassium ferrocyanide K_4_[Fe(CN)_6_], and 2 mM X-Gluc in dimethylformamide).

The plate was placed in a desiccator connected to a vacuum pump and operated in 5 cycles, consisting of 10 min of vacuum generation followed by 3 min of incubation without vacuum. This procedure was repeated until the plant material or individual organs freely settled to the bottom during incubation without vacuum. The tightly sealed plate was then incubated overnight at 37°C in the dark with shaking at 150 rpm. The following day, the staining buffer was removed, and samples were placed in absolute ethanol and incubated at 37°C with shaking, with the solution replaced every 30 minutes until chlorophyll was completely removed and the tissues became transparent. Stained samples were observed using a Zeiss SteREO Lumar.V12 and Zeiss AxioScope 2 Plus with ZEN 2.6 software.

For the sectioning, samples were embedded in LR White Acrylic Resin. Samples were incubated for 24 h in solutions with gradually increasing resin concentrations. The process began with a solution of 3 parts resin and 1 part absolute ethanol. After 12 h, samples were transferred to a 1:1 resin/absolute ethanol solution, and after another 12 h, to a 3:1 resin/ absolute ethanol solution. Subsequently, samples were incubated in resin containing a catalyst (1.88 g/100 g resin) for 24 h, replacing the resin solution halfway through. Finally, samples were placed in gelatin capsules, covered with resin and catalyst solution, sealed, and polymerised at 55°C for 24 h. Embedded samples were sectioned with a glass knife on a microtome and observed under a Zeiss AxioScope 2 Plus microscope.

### In silico analysis of transcriptomic datasets

In silico analyses of gene expression patterns were performed using publicly available transcriptomic datasets obtained from the Plant Public RNA-seq Database (PPRD; https://plantrnadb.com/) [16] and the NCBI Gene Expression Omnibus (GEO; https://www.ncbi.nlm.nih.gov/geo/). All transcriptomic projects included in this study are listed in **Table 1**.

**Table 1.** Transcriptomic datasets analyzed in this study. List of all transcriptomic BioProjects, indicating the tissue or biological process, experimental strategy (microarray or RNA-seq), BioProject/GEO accession number (in parentheses), the official project title as registered in NCBI, and the corresponding reference.

| Tissue or process | Strategy | Bioproject accession and name (NCBI) | Citation |
| --- | --- | --- | --- |
| root cap | microarray | (PRJNA97071/GSE5749) A gene expression map of the Arabidopsis root | [20] |
| root cap | microarray | (PRJNA267858/GSE63474) Gene Expression in whole roots of the Arabidopsis nlp7-1 mutant | [21] |
| root hairs | microarray | (PRJNA168641/GSE38486) Transcriptional Profiling of Arabidopsis Root Hairs and Pollen Defines an Apical Growth Signature | [22] |
| leaf petiole | RNA-seq | (PRJNA314076) High-resolution transcriptomic development map of Arabidopsis thaliana based on RNA-seq | [23] |
| leaf xylem | microarray | (PRJNA262720/GSE61941) Time course microarray analysis during xylem cell differentiation in the leaf-disk culture system | [24] |
| root xylem | RNA-seq | (PRJNA383997/GSE98097) High-resolution gene expression datasets of ontogenetic zones in the root apical meristem | [25] |
| tapetum | RNA-seq | (PRJNA254091) Transcriptomes of dyl1-3 and related bHLH mutants | [26] |
| tapetum | RNA-seq | (PRJNA274726) Transcriptome of bhlh010, bhlh089 and bhlh091 mutant anthers | [27] |
| tapetum | RNA-seq | (PRJNA397413) The tapetal cells of Arabidopsis inflorescences at stages 6-7 and 8-10 captured by laser microdissection and pressure catapulting | [28] |
| siliqua pod | RNA-seq | (PRJNA314076) High resolution transcriptomic development map of Arabidopsis thaliana based on RNA-seq | [23] |
| guard cells | microarray | (PRJNA253729/GSE58857) A transcriptional map following the developmental trajectory of the Arabidopsis stomatal lineage | [29] |
| guard cells | RNA-seq | (PRJNA484636/GSE118137) Cell-type specific transcriptome and histone modification dynamics during in situ cellular reprogramming in the Arabidopsis stomatal lineage | [30] |
| Trichome | microarray | (PRJNA112417/GSE14053) Expression data from trichomes and pavement cells | [31] |
| *V. vulnificus* inf. | RNA-seq | (PRJNA261035/GSE61418) Gene expression profiling study by RNA-seq in Arabidopsis thaliana infected by *Vibrio vulnificus* | [32] |
| *A.brassicicola* inf. | RNA-seq | (PRJNA326102/GSE83478) *Alternaria brassicicola* infection regulates miRNAs in Arabidopsis thaliana | unpublished |
| *F.oxysporum* inf. | RNA-seq | (PRJNA350767) RNAseq analysis of med18 and med20 roots infected with *Fusarium oxysporum* | [33] |
| TCV inf. | RNA-seq | (PRJNA336058/GSE85067) Comparative transcriptome analysis of TCV-infected wildtype and dcl1-9 mutant Arabidopsis thaliana | [34] |
| endoredupli. | RNA-seq | (PRJNA551314) Arabidopsis thaliana Raw sequence reads | [35] |
| endoredupli. | RNA-seq | (PRJNA754597/GSE182115) Next Generation Sequencing Facilitates Quantitative Analysis of wild-type (WT) and ERF4 transgenic plants (OE) Transcriptomes | [36] |
| endoredupli. | RNA-seq | (PRJNA812776/GSE197898) Analyses of the transcriptional and alternative splicing changes in mdf-1 in comparison to WT via RNA sequencing | [37] |
| endoredupli. | RNA-seq | (PRJNA312524) Transcriptomic effects of the cell cycle regulator LGO in Arabidopsis sepals | [38] |
| endoredupli. | microarray | (PRJNA120927/GSE19261) Photoreceptor control of shoot meristem activity and leaf initiation | [39] |

RNA-seq expression data were retrieved from the Arabidopsis RNA-seq Database, a component of PPRD (https://plantrnadb.com/athrdb/), which integrates high-throughput RNA sequencing (RNA-seq) datasets from multiple independent studies. Gene expression levels were obtained as FPKM (Fragments Per Kilobase of transcript per Million mapped reads) values provided directly by the database. No additional normalization or reprocessing of RNA-seq data was performed.

Microarray datasets were identified and selected using the NCBI Gene Expression Omnibus (GEO) repository. Only processed and normalized expression data <u>provided by the original authors</u> were used in this study. When applicable, expression values were accessed via the GEO2R web-based analysis tool (https://www.ncbi.nlm.nih.gov/geo/geo2r/) using default settings. The values used for further analysis represent normalized probe signal intensities, and no additional normalization was applied.

For each selected experiment, expression values from the available biological replicates (typically two to four, depending on the dataset) were collected and exported as CSV files. Data processing, statistical analysis, and visualization were performed using a custom R script employing the tidyverse and ggplot2 packages. Mean expression levels and standard error of the mean (SE) were calculated for each condition. Error bars presented in the figures correspond to the standard error (SE). Individual replicate values are overlaid on each bar as data points.

Statistical comparisons were performed depending on the experimental design: the Jonckheere–Terpstra test (J-T) implemented via the clinfun package, 10,000 permutations were applied to datasets with ordered conditions from developmental or infection time-course experiments; the Wilcoxon rank-sum test was used for pairwise comparisons against a wild-type control when at least one group had only two replicates, and one-way ANOVA was applied for multi-group comparisons with adequate sample sizes. Significance thresholds were set at p < 0.05 (*), p < 0.01 (**), and p < 0.001 (***). For experiments with insufficient sample size (<2 or <3, depending on the experimental design and the appropriate statistical test), values were presented only as graphs.

The sizes of the original graphs and the font sizes of their labels were adjusted in CorelDRAW to match the final dimensions of the individual panels.

### Analysis of *AtCAN1* and *AtCAN2* expression in single-nucleus RNA-seq atlas data

To examine the cell-type-resolved expression patterns of *AtCAN1* and *AtCAN2* across *Arabidopsis* development, we utilized publicly available single-nucleus RNA-seq data from the Arabidopsis Developmental Atlas [17]. Processed datasets in .h5ad format were downloaded for six developmental stages: seedling (6 days), seedling (12 days), rosette (21 days), rosette (30 days), flower, and silique, directly from the atlas portal (http://neomorph.salk.edu:9000/). Data were loaded using Scanpy [18] in Python. For each dataset, cell types annotated as “unknown” were excluded globally; “dividing” cells were additionally excluded from the seedling 6-day and rosette 21-day datasets. For the silique dataset, embryo-associated cell types and the “younger silique stress” cluster were excluded from the analysis.

For each developmental stage, expression matrices were extracted for *AtCAN1* and *AtCAN2* and organized by annotated cell type. Two summary statistics were computed per gene–cell type combination. First, the percentage of expressing nuclei was calculated as the number of nuclei with a normalized count strictly greater than zero divided by the total number of nuclei in that cell type. Second, the mean normalized expression was calculated as the arithmetic mean of normalized counts across all nuclei in the cell type, including nuclei with zero counts. These metrics were computed using the expression values as stored in the atlas .h5ad files, which represent normalized (but not log-transformed) counts as processed by the original authors. No additional normalization or transformation was applied. Results were organized into per-stage summary tables and visualized as heat maps using seaborn, with color intensity reflecting the percentage of expressing nuclei (viridis color scale, independently normalized per developmental stage to its maximum observed value) and numerical annotations indicating the mean normalized expression value.

The sizes of the original graphs and the font sizes of their labels were adjusted in CorelDRAW to match the final dimensions of the individual panels.

### Source and Genotypic Verification of Arabidopsis thaliana Mutant Line

Arabidopsis thaliana ecotype Columbia (Col-0) was used as the wild type. The insertion mutant line GABI_022B01 (SALK) was obtained from the Nottingham Arabidopsis Stock Centre (NASC) as line CS402029. The presence of the wild-type allele was confirmed by PCR using the LP primer (TCCCACTGAATTCAAAACACC) in combination with the RP primer (GAAGCCTCCAGGCTTGTTATC). The presence of the T-DNA insertion in the AtCAN1 gene (At3g56170) was verified using the RP primer together with the T-DNA BP primer (ATAATAACGCTGCGGACATCTACATTTT). Homozygous mutants were selected in the F3 generation and confirmed by PCR genotyping. The absence of the full-length AtCAN1 transcript in the homozygous line was confirmed by RT-PCR (data not shown).

Arabidopsis plants were grown in 50-mL pots containing sterile soil in a phytotron under a 16-h light/8-h dark photoperiod. Day and night temperatures were maintained at 22°C and 18°C, respectively. Plants were watered and fertilized daily with MS mineral solution.

### Rosette leaf area measurement

At 30 days after germination (DAG), rosettes were photographed from above on a dark background alongside a red calibration square (4 cm^2^). Rosette leaf area was measured from digital images using the Easy Leaf Area software [19]. Statistical analysis was performed using Welch’s t-test in R. For figure preparation, the red calibration square was masked from representative images. No other image manipulations were performed. Original unmodified images are available upon request.

## Results

### Production of *AtCAN1* promoter-GUS and *AtCAN2* promoter-GUS transgenic plants

The analysis of expression profiles is a basic approach to understanding biological function. In the studies presented below, we assumed that identifying the expression of SNc family genes in tissues undergoing well-defined nucleic acid degradation processes would support the hypothesis that these genes participate in such processes. This strategy was successfully applied in previous research, where analysis of plants transformed with a chimeric construct (*BFN1* promoter-GUS) demonstrated the involvement of the BFN1 (ENDO1) nuclease from the S1/P1 family in plant PCD [10]. In this work, we used a similar methodology. We selected potential promoter sequences of the two *A. thaliana* SNc nucleases (2899 bp for *AtCAN1* and 1814 bp for *AtCAN2*) and introduced them into the modified pCAMBIA3301 genetic vector upstream of the GUS reporter gene. These constructs were introduced into *Arabidopsis* plants using the floral dip method, and the resultant transgenic lines were analysed for SNc promoter activity during plant development. The activity of the GUS reporter gene under the control of the selected promoters was examined comprehensively across all organs and developmental phases; however, in this paper, we present only the results demonstrating detectable GUS protein activity.

### Expression pattern of *AtCAN1* during early stages of plant development

We began our study of the expression profile of *AtCAN1* nuclease in *A. thaliana* by detecting GUS signals at early stages of plant development, from seed germination through the third week of growth (**Figure 1**). The first appearance of the GUS signal was observed in two-day-old seedlings. At this stage, the strongest signal was detected in the root cap. In addition, a pronounced GUS signal was visible in the transition zone between the hypocotyl and the root, with a particularly high density in the root hairs. Furthermore, in this part of the plant, the GUS signal was also present along the developing axial cylinder of the root. A weaker signal was also detected in the cotyledon area; however, it disappeared during subsequent stages of seedling development.

**Figure 1.**
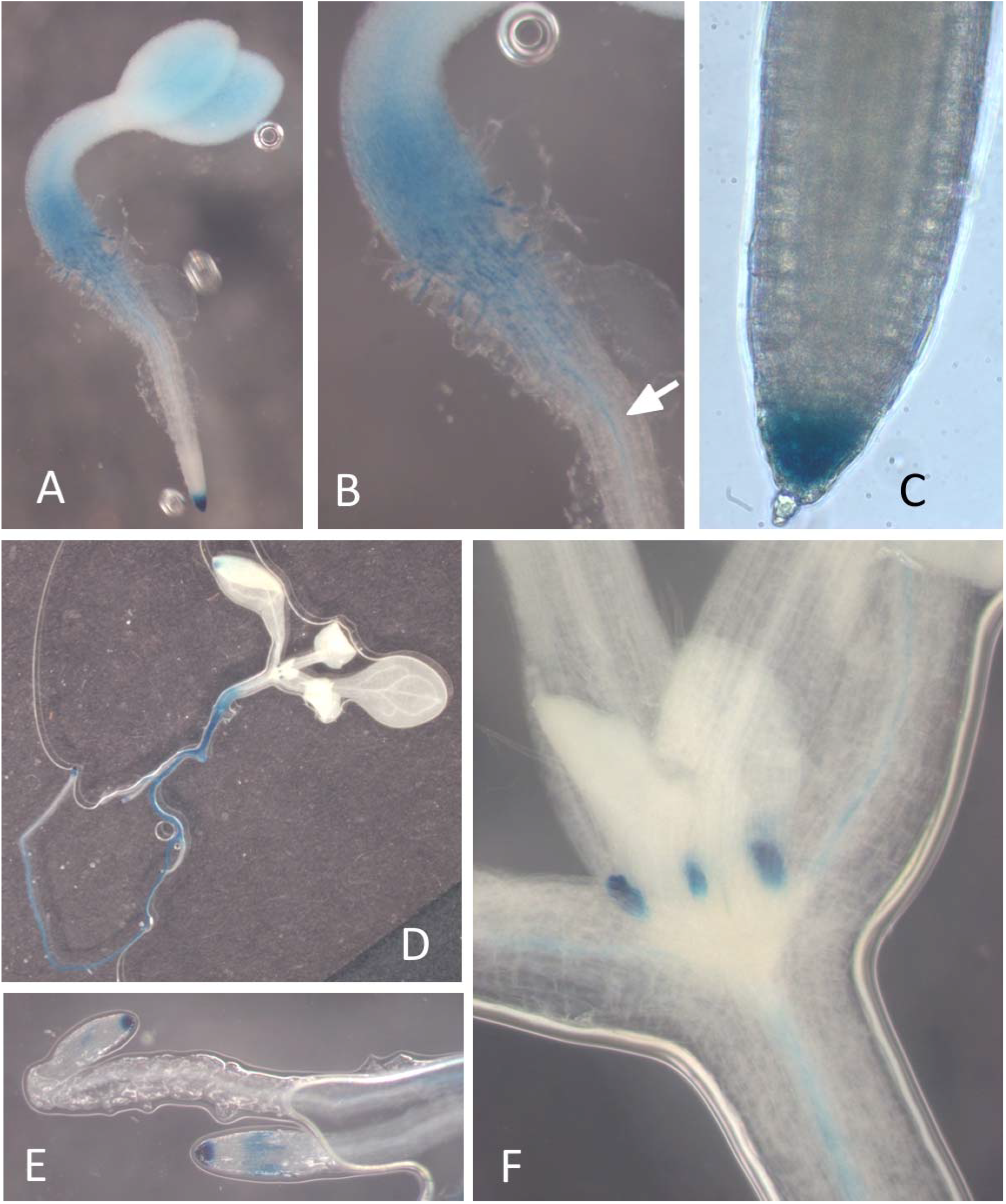
Histochemical staining of GUS activity driven by the *AtCAN1* promoter in *A*.*thaliana* seedlings. **(A)** Overall view of a two-day-old seedling. **(B)** Enlargement of the transition zone between the hypocotyl and the root, showing GUS signal in the root hairs and in the developing axial cylinder (arrow). **(C)** Enlargement of the root cap. **(D)** Overall view of a 7-day-old seedling. **(E)** Enlargement of the terminal, younger parts of a 7-day-old seedling root. **(F)** Enlargement of the floral shoot apex region, with GUS signals in the stipules and in the developing vascular bundles of stem and leaf petioles.

As shown in **Figure 1**, which presents the staining results of a 7-day-old seedling, in the following days, the GUS signal extended along the entire length of the root, except for the youngest sections, where it remained confined to the root cap. In the above-ground part of the seedling, a specific GUS signal appeared in the stipules and along the vascular bundles of the stem and leaf petioles.

### Expression pattern of *AtCAN1* in vegetative organs of mature plants

In the vegetative organs of mature plants, the activity of the *AtCAN1* promoter was detected in several locations (**Figure 2**). In 14-day-old plants, intense GUS signals were observed in the petioles of rosette leaves. Furthermore, GUS activity was also detected in the inflorescence stem and the leaf blade area. In the stem, the GUS signal persisted in the vascular bundles and on its surface, where distinct and highly specific GUS signals were observed in stomata and trichomes. We also observed *AtCAN1* promoter activity in several leaf structures. To further characterize these signals, we analyzed cotyledons, rosette leaves, and stem leaves. When analyzing rosette leaves, we included both juvenile rosette leaves with smooth margins and adult leaves with serrated margins. Analysis of all these cases showed that the GUS signal in leaves was detected in the vascular bundles, trichomes, and hydathodes. However, the intensity of individual signals varied depending on the leaf category and growth conditions–for example, whether the plants were grown in soil or *in vitro* under relatively high humidity exceeding 95%. Signals in hydathodes were observed both along the smooth leaf margin and at the tips of the leaf serrations, which are tooth-like protrusions that also contain hydathodes in *Arabidopsis* [40]. Interestingly, while the GUS signal was present in both stem and leaf trichomes, it was detected only in stem stomata and not in those of leaves.

**Figure 2.**
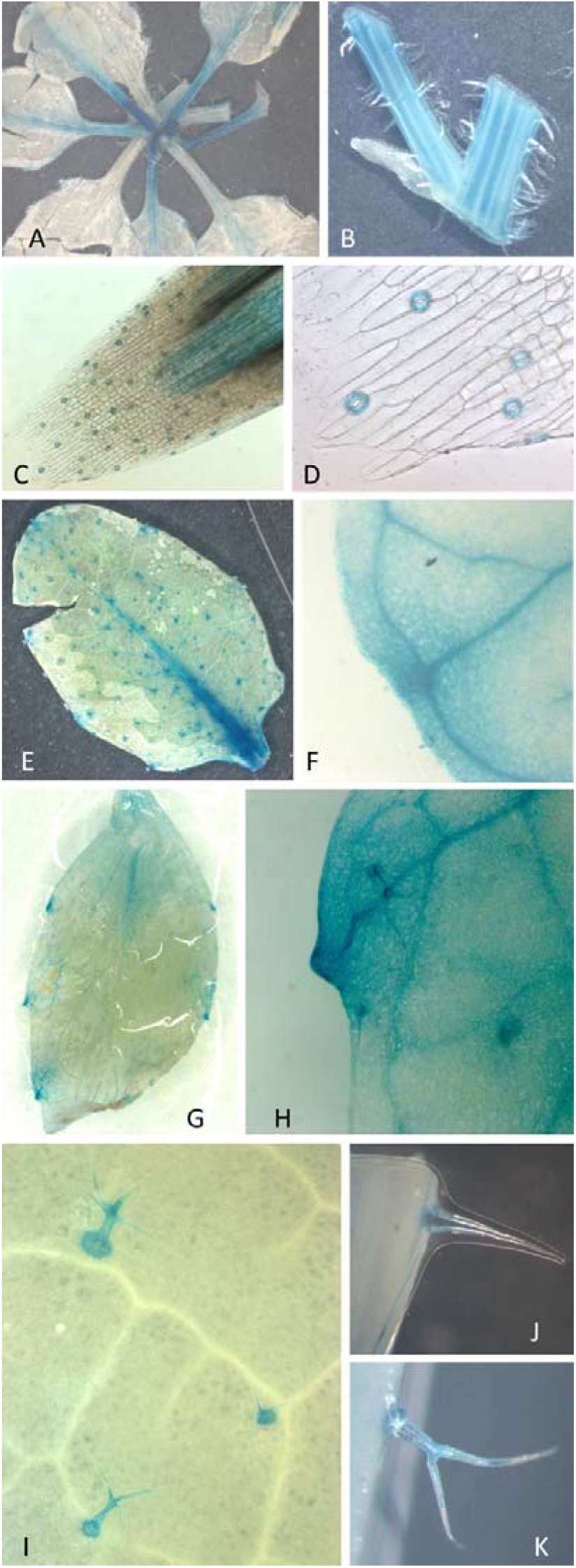
Histochemical staining of GUS activity driven by the *AtCAN1* promoter in vegetative organs of mature plants. **(A)** 21-day-old *A. thaliana* plant with dominant GUS signal in leaf petioles. **(B)** Cross-section of the lower part of the flower stem with an attached leaf petiole, showing GUS signals in the vascular bundles. **(C)** microscopic image of an oblique stem section, showing GUS signal in stomata and vascular bundles. **(D)** Enlargement of the stem epidermis with GUS signal in stomata. **(E)** Overall view of a rosette leaf. **(F)** Enlargement of a leaf region showing GUS signal in a hydathode located at the smooth leaf margin. **(G)** Overall view of a rosette leaf with serrated margins. **(H)** Enlargement of a GUS signal in a hydathode located at the tooth-like protrusion. **(I)** Trichomes on the leaf surface. **(J)** Enlargement of the stem trichome. **(K)** Enlargement of a single trichome at the edge of the leaf.

### Expression of *AtCAN1* during leaf senescence

The spatial distribution of the GUS signal in leaves showed clear alterations as senescence progressed. In fully developed, mature leaves, GUS activity was detected only in the previously mentioned, narrowly defined leaf structures. It was not detectable in the green tissues, including the mesophyll and epidermis (**Figure 2**). As illustrated in **Figure 3**, however, the pattern of GUS expression underwent a pronounced shift upon the onset of leaf wilting. The signal initially emerged in tissues displaying chlorosis at the leaf margins. Subsequently, it expanded as senescence progressed, thereby providing clear evidence of *AtCAN1* promoter activation in response to senescence-associated factors.

**Figure 3.**
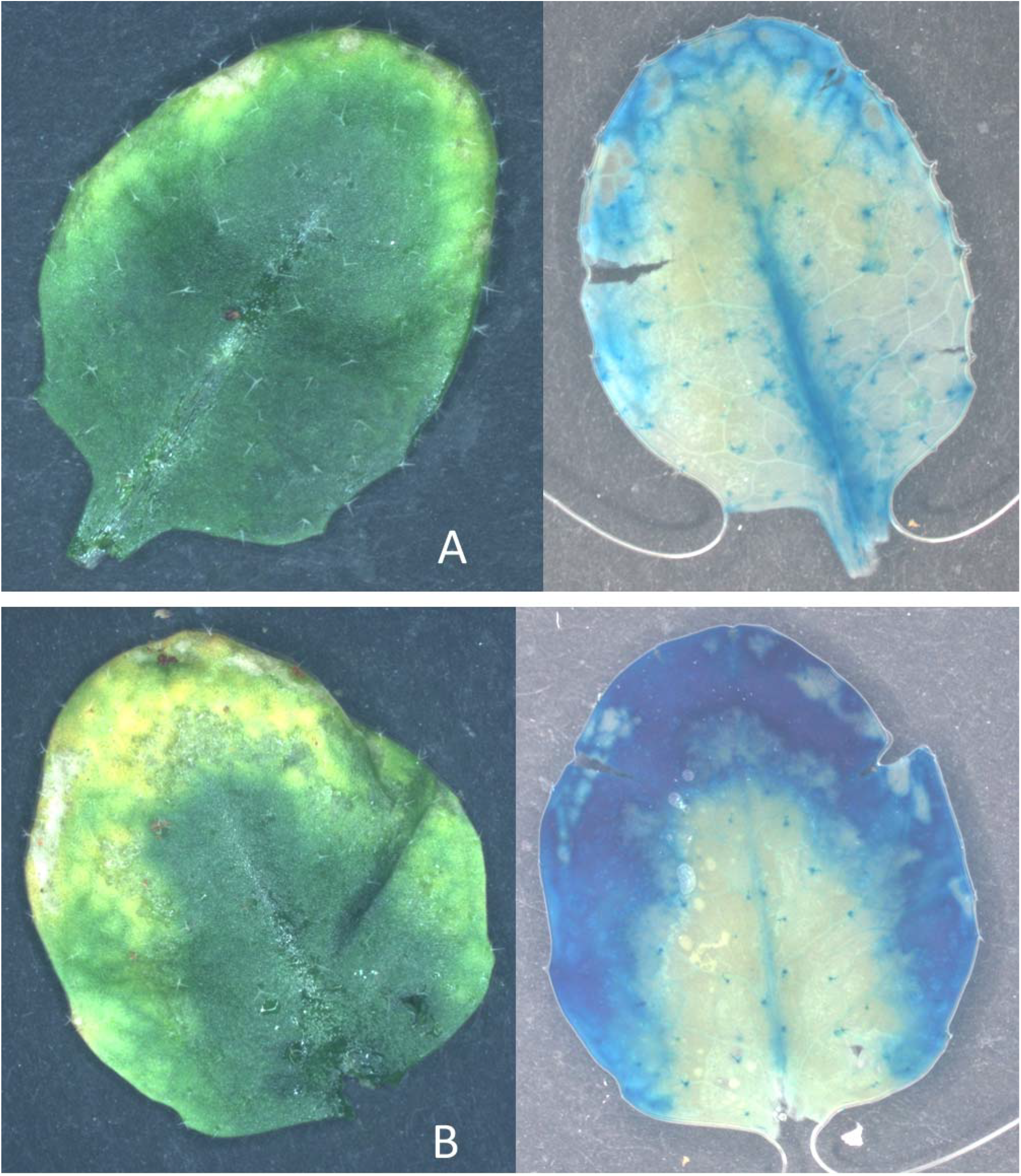
Histochemical staining of GUS activity driven by the *AtCAN1* promoter in senescing leaves. **(A)** A leaf at an early stage of senescence, before (left) and after (right) GUS staining. **(B)** A leaf at a later stage of senescence, before (left) and after (right) GUS staining.

### Spatial pattern of *AtCAN1* expression in floral tissue and developing siliques

Expression of the *AtCAN1* promoter was also observed in flowers (**Figure 4**). Between stages 10 and 16 of flower development (according to the classification proposed by [41] the GUS signal was detected exclusively in the anthers. At these stages, expression was observed both in the tapetum and the developing pollen. As flower development progressed, the distribution of the signal changed: it disappeared from the tapetum as this tissue degraded and became localized in the pollen grains. This expression pattern is characteristic of tapetum–derived proteins, which are known to be incorporated into pollen during its maturation.

**Figure 4.**
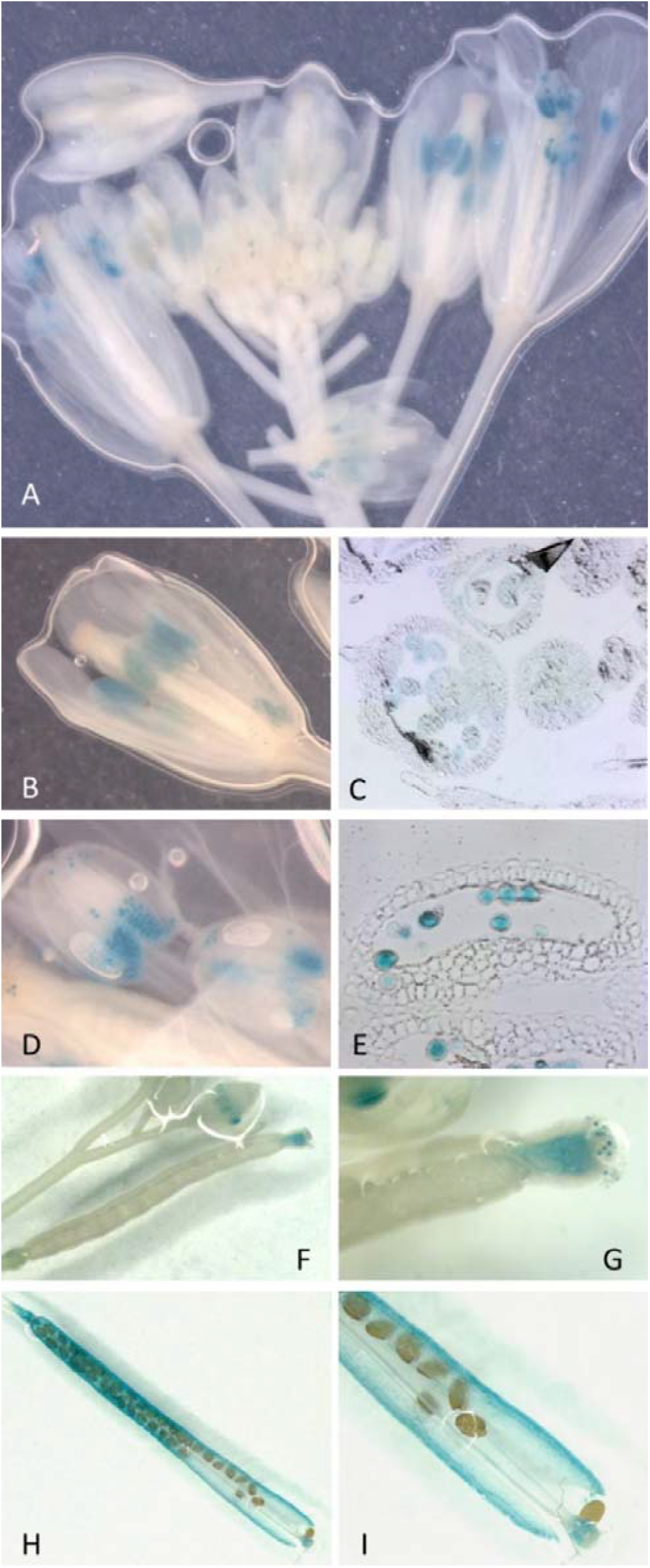
Histochemical staining of GUS activity driven by the *AtCAN1* promoter in *Arabidopsis* flowers and developing siliques. **(A)** Overall view of the inflorescence. **(B)** Enlargement of a flower at the 14th stage of development. **(C)** Microscopic image of a cross-section of anthers at the 14th stage of flower development. **(D)** Enlargement of the flower at the 15th stage of development. **(E)** Microscopic image of a cross-section of anthers at the 15th stage of flower development. **(F)** Overall view of an early silique. **(G)** Enlargement of the transmitting tissue of the pistil style. **(H)** Overall view of a late silique. **(I)** Enlargement of the late silique showing GUS signal in the silique valves.

At later stages, GUS activity was also detected during the transformation of the pistil into seed pods. Initially, a highly specific signal was observed in the transmitting tissue of the pistil style. In contrast, at later stages of silique maturation, the signal became visible along the entire surface of the valves (**Figure 4**).

### Expression of the *AtCAN2* promoter in plant tissues

Observations similar to those obtained for the gene encoding AtCAN1 were also made for the second member of the SNc family, *AtCAN2*. Despite considerable similarity in the catalytic properties of these two enzymes, their gene expression profiles differed markedly. Expression of the *AtCAN2 promoter::GUS* construct was detected in only two cases. In the seedling, *AtCAN2*, like AtCAN1, showed strong, highly specific expression in stipules. In mature plants, we detected the GUS signal only in the leaf hydathodes (**Figure 5**). No GUS signal was observed in other tissues in which *AtCAN1 promoter::GUS* activity had previously been detected.

**Figure 5.**
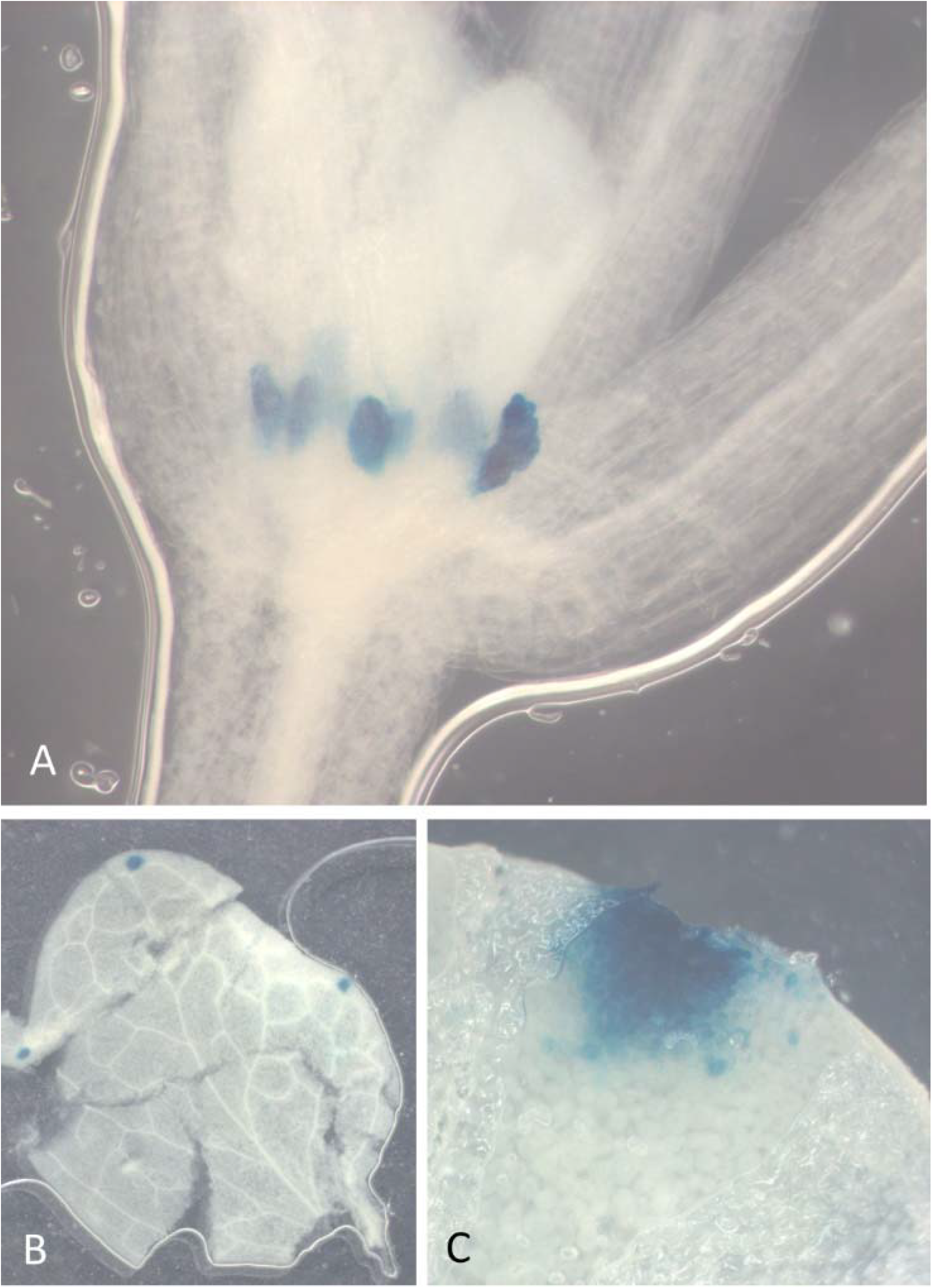
Histochemical staining of GUS activity driven by the *AtCAN2* promoter in an *Arabidopsis* plant. **(A)** The floral shoot apex region of a 7-day-old seedling, showing GUS signal in the stipules. **(B)** Overall view of a mature leaf. (**C**) Enlargement of a leaf region showing GUS signal in the hydathodes.

### Transcriptomic evidence for tissue-specific expression of *A. thaliana* SNc family genes

The results from GUS reporter assays demonstrate that SNc nucleases are expressed in distinct, well-defined plant tissues. Considering the nucleolytic activity of these enzymes, their previously reported subcellular localization, and the characteristics of the tissues in which they are expressed, these findings support hypotheses regarding their biological function. However, an important condition for this type of analysis is independent validation of the tissue-specific expression patterns observed in this study. Although the GUS reporter assay is a widely used and well-established approach, several caveats must be considered when interpreting the results, as they may affect the robustness of the conclusions. One such limitation is that the DNA fragment used as the promoter region may be incomplete, since gene expression can also be regulated by cis-acting elements located outside the analysed promoter sequence. In addition, transgene expression may in some cases be influenced by the chromatin environment at the site of genomic insertion.

For these reasons, the results obtained using the GUS reporter assay were compared with independent analyses reflecting gene expression under native conditions. To this end, a comparative analysis was performed between the GUS-based expression patterns presented above and expression data derived from genome-wide transcriptomic profiling approaches, primarily microarray- and RNA-seq–based assays (**Table 1**). Because multiple research groups have independently analysed the same transcriptomes, these datasets provide a robust and objective reference for assessing gene expression patterns. Moreover, the available transcriptomic data include analyses of numerous plant mutants, providing additional insights into the mechanisms underlying the regulation of the analysed gene’s expression.

The root cap is one of the plant tissues in which the GUS reporter signal driven by the *AtCAN1* promoter was particularly strong and highly localized. Transcriptome profiling data further support this observation. For example, a comparative analysis of root transcriptomes from the lateral root cap (LRC) and the adjacent elongation zone (EZ) [20], as shown in **Figure 6A**, revealed a clear enrichment of *AtCAN1* expression in the LRC relative to the EZ. In contrast, no such enrichment was observed for *AtCAN2*, consistent with our previous results, which did not indicate elevated expression of this gene in the LRC relative to other root regions.

**Figure 6.**
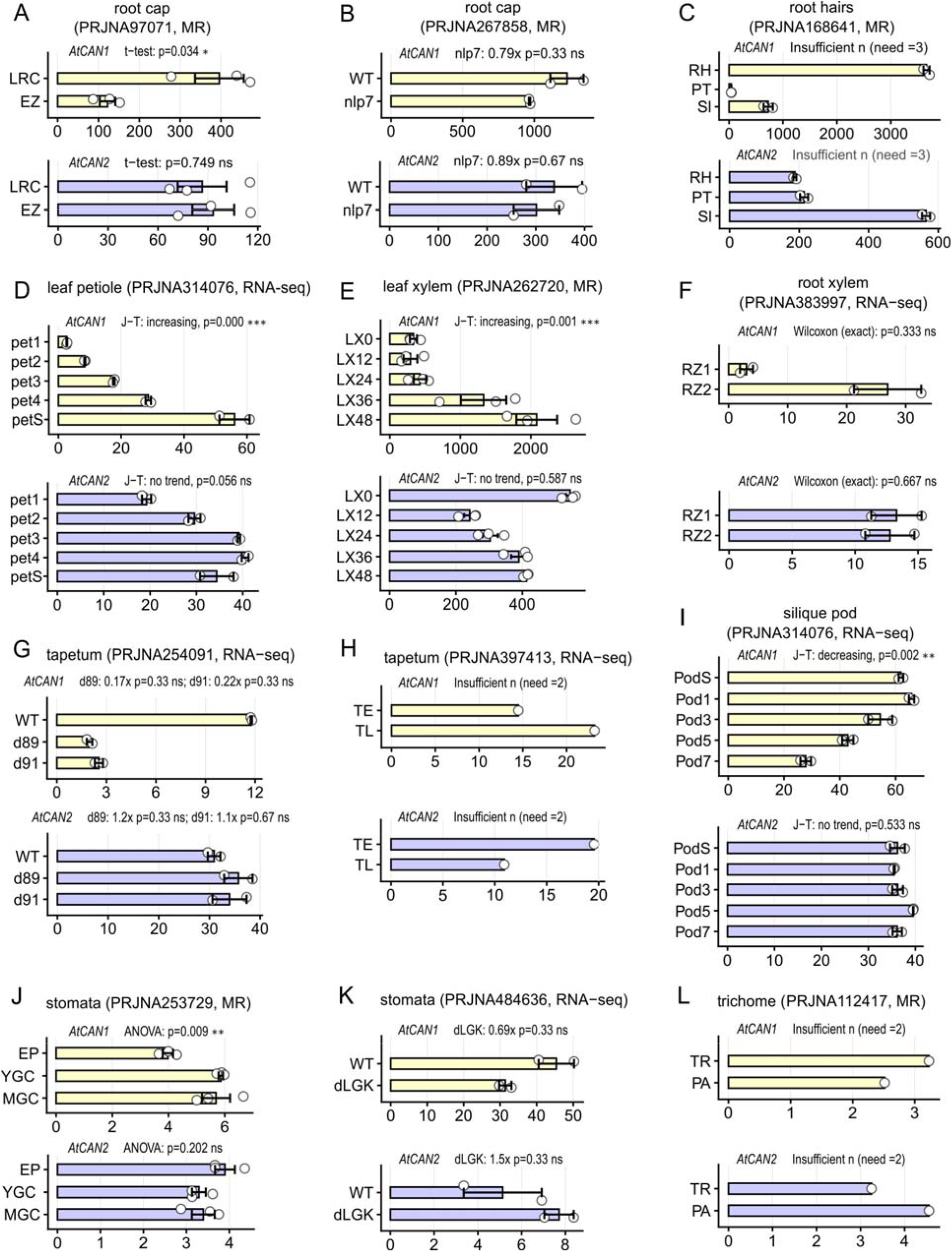
Transcriptome-based analysis of *AtCAN1* and *AtCAN2* gene expression across different *A. thaliana* tissues or in response to various experimental treatments. In each panel, two graphs show the expression of the genes encoding AtCAN1 (above, yellow) and AtCAN2 (below, purple). Expression data were retrieved and processed as described in the Methods section. BioProject accession numbers as listed in Table 1 and the type of experiment, i.e., RNA sequencing (RNA-seq) or microarray (MR), are given in parentheses. Gene expression levels are presented as FPKM for RNA sequencing or normalized expression values for microarray datasets (see Materials and Methods). Detailed information about individual experiments is provided in the Results section. Statistical annotations above each graph indicate the test used and the result: J–T, Jonckheere–Terpstra trend test (with direction: increasing, decreasing, or no trend); t-test, Student’s t-test; ANOVA, one-way ANOVA; Wilcoxon (exact), Wilcoxon rank-sum test. For pairwise comparisons against wild-type controls with limited replicates, fold-change relative to WT and the Wilcoxon p-value are shown (e.g., “0.17x p=0.33”). Significance levels: *p < 0.05, **p < 0.01, ***p < 0.001; ns, not significant. “Insufficient n” indicates that the number of replicates was too low for the appropriate statistical test. **(A)** Experiment demonstrating the expression of the analyzed genes in the lateral root cap (LRC) compared to the adjacent root elongation zone (EZ). **(B)** Effect of the dNLP7 mutation on the expression of the analyzed genes. (**C**) Experiment showing different expression levels of the analyzed genes in two cell types representing a similar mechanism of apical growth, i.e., root hairs (RH) and pollen tubes (PT), as well as in siliques (SI). (**D**) Expression levels of the studied nuclease genes in the petioles of young (Pet1), intermediate (Pet2, Pet3), mature (Pet4), and senescent (PetS) leaves. (**E**) Expression of the analyzed genes in leaf-disk cultures induced for xylogenesis at time points from 0 h (LX0) to 48 h (LX48). (**F**) Expression of the studied nuclease genes in the root xylem maturation zone (RZ2) compared with the region located below (RZ1). (**G**) Expression of the analyzed genes in *A. thaliana* bhlh89 (d89) and bhlh91 (d91) mutants with impaired tapetum development. (**H**) The level of expression of the studied genes in tapetal cells at early (TE) and late (TL) stages of development, leading to programmed cell death (PCD). (**I**) The level of expression of the analyzed genes in five silique pods at different stages of ageing, from Pod7 (youngest) to Pod1 (oldest) and PodS (senescing). (**J**) The level of expression of the studied genes at different stages of stomatal development, namely epidermal cells (EP), young guard cells (YGC), and mature guard cells (MGC). (**K**) Reduced expression of the analyzed genes in *A. thaliana* FAMA^LGK^ mutants (dLGK) with impaired guard cell identity. (**L**) Comparison of the expression levels of the analyzed nuclease genes in trichomes (TR) and pavement epidermal cells (PA).

Additional evidence for the association of AtCAN1 with the root cap is provided by comparative transcriptome analysis of wild-type Arabidopsis (WT) and a mutant lacking functional NLP7 protein. NLP7 (NIN-LIKE PROTEIN 7) is a transcription factor that specifically regulates root cap maturation in Arabidopsis and is highly expressed in the columella root cap. Previous studies have shown that loss of NLP7 function severely disrupts the development of this tissue [21]. As shown in **Figure 6B**, the dNLP7 mutant also exhibits a marked reduction in AtCAN1 expression, further supporting a functional association of this nuclease with the root cap.

Another root region in which our analyses revealed strong AtCAN1 promoter activity was the root hair zone. This observation was further supported by bioinformatic analysis of transcriptomic datasets. One illustrative example comes from studies comparing the transcriptomes of root hairs and pollen tubes, based on the premise that these cell types share similarities related to an analogous mechanism of apical growth. In that study, the authors also compared these data with transcriptomes from six additional vegetative organs of *Arabidopsis thaliana* — seedling, flower, ovule, unpollinated pistil, silique, and leaf [22]. Our analysis of these datasets demonstrated that, among the eight tissues examined, AtCAN1 expression was highest in root hairs, markedly exceeding the levels observed in pollen tubes. The remaining tissues included in this experiment displayed substantially lower AtCAN1 expression, with the highest levels detected in siliques (**Figure 6C**). Notably, the expression profile of AtCAN2 deviated from this pattern.

In our study, GUS activity driven by the *AtCAN1* promoter was also detected in the vascular bundle region. This finding supports our previous hypothesis linking AtCAN1 expression to xylem formation, proposed in Lesniewicz *et al*. [12], which describes the catalytic activity and subcellular localization of this nuclease. That hypothesis was based on microarray data from the Genevestigator database, whereas the transcriptomic analyses presented here provide further, more detailed evidence supporting this suggestion. The research project “High-resolution transcriptomic developmental map of Arabidopsis thaliana based on RNA-seq” (**Table 1**) [23] provided extensive data on transcriptomic changes associated with different developmental stages of individual organs, including the leaf petiole. As shown in **Figure 6D**, these data confirm our observations, indicating a strong increase in AtCAN1 expression in this organ.

Additional transcriptomic profiling enabled a more detailed analysis of *AtCAN1* expression in individual components of the vascular bundle [24, 25]. The experiment employing an in vitro leaf-disk culture system to induce xylogenesis demonstrated that *AtCAN1* expression increases during leaf xylem development (**Figure 6E**). In turn, an experiment employing FACS coupled with RNA sequencing to isolate and analyze xylem-specific cell populations from distinct ontogenetic zones of the root revealed that the level of *AtCAN1* expression increases in the root xylem maturation zone compared with the region located between this zone and the merystem (**Figure 6F**). These data also indicate that the expression pattern of the gene encoding AtCAN2 differs from that of AtCAN1 and does not show a clear association with xylogenesis.

The above data on *AtCAN1* expression in the xylem and the root cap suggest that this gene may be involved in different types of programmed cell death (PCD). Therefore, we considered it particularly important to obtain independent confirmation of our GUS-based results suggesting an association between AtCAN1 and tapetum degradation, as this tissue is widely regarded as a model system for developmental PCD. For this reason, the results of transcriptome profiling studies examining the impact of mutations in the genes encoding the bHLH89 and bHLH91 proteins on tapetum development were considered particularly informative. These proteins are basic helix–loop–helix (bHLH) transcription factors that specifically regulate tapetum and pollen development. Deletion or disruption of these genes in Arabidopsis has been shown to cause severe defects in tapetum development [26]. As shown in **Figure 6G**, analysis of RNA-seq–derived transcriptomes from *bhlh89* and *bhlh91* mutants revealed that these mutations significantly reduce AtCAN1 expression. Similar results, supporting an association between *AtCAN1* expression and tapetum development, were obtained from an independent transcriptomic study by Zhu et al. [27], which also investigated the effects of bhlh089 and bhlh091 on tapetum development (data not shown). Further evidence for the involvement of AtCAN1 in PCD is provided by a comparison of transcriptomes from tapetal cells at stages 6–7 and 8–10, captured by laser microdissection and pressure catapulting [28]. As shown in **Figure 6H**, AtCAN1 expression is markedly higher at stages 8–10, which correspond to tapetum degradation via PCD, than at earlier stages.

Another plant structure in which the GUS signal correlated with PCD progression was the silique wall. This tissue was also included among the samples analysed within the previously mentioned project PRJNA314076 [23] (**Table 1**). As shown in **Figure 6I**, seedless silique pods subjected to transcriptomic analysis at different stages of ageing exhibited a pronounced increase in AtCAN1 expression levels. In contrast, the expression of AtCAN2 did not show a similar trend, consistent with the previously analysed cases.

AtCAN1 promoter activity was detected not only in tissues undergoing developmental PCD but also in tissues not associated with this process, as evidenced by the GUS signal observed in guard cells. Multiple transcriptomic datasets support this observation. Specifically, transcriptomic analyses of stomatal development stages [29] demonstrated elevated AtCAN1 expression in guard mother cells and mature guard cells compared with epidermal cells (**Figure 6J**).

The involvement of AtCAN1 in guard cell development is further supported by studies characterizing the cell– type–specific transcriptome of Arabidopsis stomatal lineages [30]. In this project, the authors compared transcriptomes of wild-type (WT) plants with those of FAMA^LGK^ mutants. FAMA is a basic helix–loop–helix transcription factor that is both necessary and sufficient for a cell to acquire and maintain stomatal guard cell (GC) identity. In contrast, point mutations in FAMA (LGK) fail to sustain the terminal identity of GCs. Comparative transcriptomic analysis of WT and LGK lineages revealed that AtCAN1 expression is reduced in the mutant background (**Figure 6K**).

Stomata are among the epidermal sites most susceptible to microbial infection. Therefore, given the potential role of SNc nucleases in guard cells and other cells highly exposed to infection, such as root hairs and hydathodes, we explored whether these enzymes contribute to plant responses to biotic stress. Such a role was hypothesized in our previous work [12], based on microarray data indicating elevated SNc gene expression following *Pseudomonas* infection. Analysis of currently available microarray and RNA-seq datasets supports this hypothesis. In addition to further evidence linking AtCAN1 expression to *Pseudomonas* infection, we observed increased expression of this gene in response to infection by other bacterial species, such as *Vibrio vulnificus* [32] as well as by fungi, including *Alternaria brassicicola* and *Fusarium oxysporum* [33], and by Turnip crinkle virus - TCV [34] (**Figure 7A–D**). Notably, the induction of *AtCAN1* expression in plants infected by *F. oxysporum* was significantly stronger in Arabidopsis *med25* and *med8* mutants than in WT plants. Because mutations in MED18 and MED20 result in downregulation of jasmonate-associated genes and upregulation of salicylic acid–associated pathogenesis-related genes [33], these findings suggest that *AtCAN1* is regulated predominantly via the salicylic acid signaling pathway. It is worth noting that although *AtCAN1* and *AtCAN2* displayed distinct expression patterns in most previous analyses, both genes responded similarly to the biotic stresses examined here.

**Figure 7.**
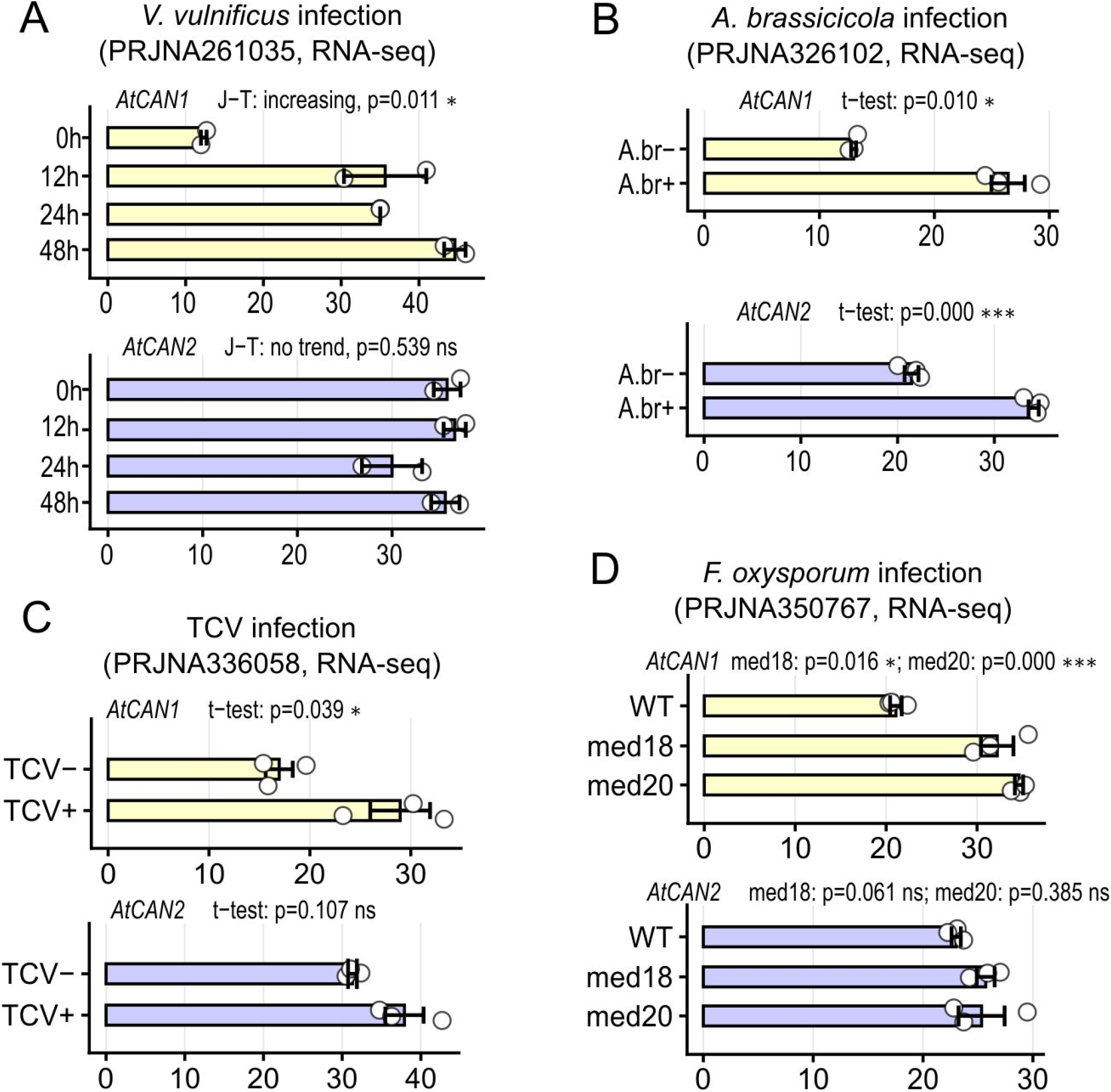
Transcriptome-based analysis of AtCAN1 and AtCAN2 gene expression in response to various pathogenesis-related stimuli. In each panel, two graphs show the expression of the genes encoding AtCAN1 (above, yellow) and AtCAN2 (below, purple). All methodological and technical details, including statistical test abbreviations, are as described in Figure 6. (**A**) Expression levels of the studied genes from 0 (0h) to 48 h (48h) post-infection with *Vibrio vulnificus*. (**B**) Expression levels of the studied genes in plants without (A.br−) and with (A.br+) infection by the pathogenic fungus *Alternaria brassicicola*. (**C**) Expression levels of the studied genes in plants without (TCV−) and with (TCV+) infection by Turnip crinkle virus (TCV). (**D**) Expression levels of the studied genes in WT and in MED18 (med18) or MED20 (med20) mutant plants upon infection with the pathogenic fungus *Fusarium oxysporum*.

One of the most notable findings of this study was the detection of AtCAN1 promoter activity in structures that do not appear to be directly associated with programmed cell death (PCD), such as trichomes and stipules. Such small or transient plant structures are rarely included in comprehensive transcriptomic analyses. Nevertheless, evidence for high AtCAN1 expression in trichomes was identified. This is illustrated in **Figure 6L**, which is based on data from a comparative transcriptomic analysis of cell sap derived from trichomes and pavement epidermal cells [31]. Elevated expression of AtCAN1 in trichomes is further supported by the results reported by Jakoby et al. [42], whose transcriptomic analysis demonstrated that the gene encoding AtCAN1 ranks among the top 5% of the most highly expressed genes in mature Arabidopsis trichomes.

Because a shared characteristic of Arabidopsis structures, such as trichomes and stipules, is endoreduplication, we examined whether SNc nuclease expression correlates with this process. Analysis of multiple transcriptomic datasets derived from experiments in which endoreduplication was induced revealed that this process was frequently associated with increased expression of one or both SNc nucleases. Consistent with these observations, analysis of transcriptomic data from studies of the Arabidopsis circadian clock [35] showed that elevated AtCAN1 expression accompanies endoreduplication resulting from cell-cycle alterations during differentiation induction in the VISUAL system (**Figure 8A**).

**Figure 8.**
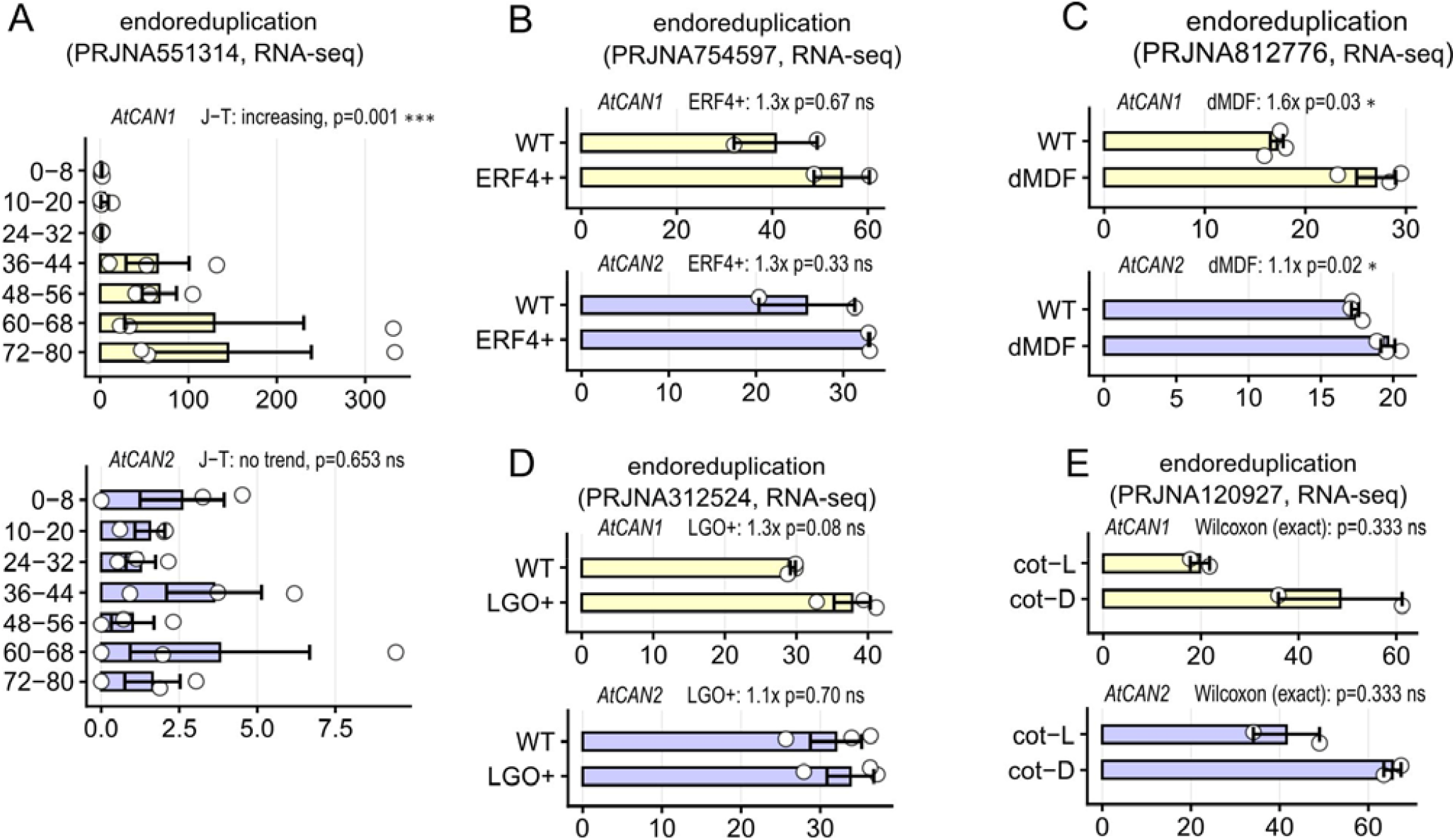
Transcriptome-based analysis of AtCAN1 and AtCAN2 gene expression in *A. thaliana* cells with induced endoreduplication. In each panel, two graphs show the expression of the genes encoding AtCAN1 (above, yellow) and AtCAN2 (below, purple). All methodological and technical details, including statistical test abbreviations, are as described in Figure 6. (**A**) Correlation between *AtCAN1* and *AtCAN2* gene expression levels and the endoreduplication process occurring as a result of differentiation induction in the VISUAL system. Time intervals (from 0–8 to 72–80 h) denote the periods following initiation of endoreduplication. (**B**) Expression levels of the analyzed genes in wild-type (WT) plants and in plants with elevated endoreduplication induced by ERF4 overexpression (ERF4+). (**C**) Expression levels of the nuclease genes in loss-of-function MDF1 mutants (dMDF) compared with wild-type (WT) plants. (**D**) Expression levels of the analyzed genes in wild-type (WT) plants and plants with increased endoreduplication induced by LGO overexpression (LGO+). (**E**) Expression levels of the analyzed genes in cotyledons induced to undergo endoreduplication through growth in darkness (cot-D) relative to light-grown control cotyledons (cot-L).

Consistent with this relationship, increased *AtCAN1* expression was also observed in response to genetic modifications that promote elevated endoreduplication levels. One such example is the overexpression of ETHYLENE-RESPONSIVE ELEMENT BINDING FACTOR 4 (ERF4) (**Figure 8B**). ERF4 functions as a positive regulator of endoreduplication by acting as a transcriptional repressor of cell cycle–related genes. Plants overexpressing ERF4 have been shown to regulate cell size by modulating nuclear endopolyploidy levels [36].

A similar effect on *AtCAN1* expression was observed in studies analysing mdf1 loss-of-function mutants (**Figure 8C**). MDF1 is a conserved splicing factor involved in the downregulation of mitotic regulators, thereby modulating cell cycle progression. Loss of MDF1 function is associated with growth defects and impaired entry into mitosis, which in turn leads to increased endoreduplication [37].

The extent of endoreduplication is also influenced by LGO (LOSS OF GIANT CELLS FROM ORGANS), which acts as an inhibitor of cyclin-dependent kinases (CDKs). Schwarz and Roeder [38] demonstrated that LGO functions as a key transcriptional switch, directing specific gene expression programs during sepal development. Their study showed that endogenous LGO regulates the transition to endoreduplication, thereby controlling the balance between giant and small cells. Transcriptomic analyses of this developmental process further revealed that LGO overexpression in sepals promotes the formation of endoreduplicated giant cells. Consistent with this, transcriptomic datasets from this study indicate that increased endoreduplication is accompanied by elevated AtCAN1 expression (**Figure 8D**).

Evidence for a correlation between *AtCAN1* expression and endoreduplication is also provided by experimental studies in which this process was induced by environmental conditions [39]. This relationship was demonstrated in an experiment showing increased levels of endoreduplication in cotyledon cells maintained under dark conditions, compared with the control consisting of seedlings grown under light conditions (**Figure 8E**).

To independently validate our SNc genes expression results, we also used data from the *Arabidopsis* Developmental Atlas Viewer [17]. This resource provides comprehensive transcriptomic coverage across the entire life cycle of *Arabidopsis thaliana*. The atlas is based on large-scale single-nucleus RNA sequencing (snRNA-seq) of whole tissues sampled at multiple developmental stages, including seeds, seedlings, rosettes, stems, flowers, and siliques. Transcriptomes from over 400,000 quality-filtered nuclei were clustered into distinct cell types using unsupervised clustering of snRNA-seq expression profiles. This database enables systematic comparison of gene expression levels across cell types and analysis of expression dynamics across developmental stages within a given *Arabidopsis* organ. Moreover, an important advantage of these data is that they allow direct comparison of expression patterns of different genes within the same tissue or cell type.

Transcriptomic analysis of six-day-old seedlings revealed relatively high *AtCAN1* expression among cells annotated as vascular. It should be noted that, because mature xylem cells are dead and mature phloem cells are transcriptionally inactive, the term *vascular* refers here exclusively to early developing xylem and phloem cells, as well as to vascular cell types that retain metabolic activity, such as procambium, cambium, and companion cells. In six-day-old seedlings, *AtCAN1* expression was also markedly higher in trichomes and trichoblasts compared with the other samples (**Figure 9A**). Consistent results were obtained from analyses of 12-day-old seedlings, in which the highest AtCAN1 expression level was likewise observed in xylem and trichoblasts (**Figure 9B**).

**Figure 9.**
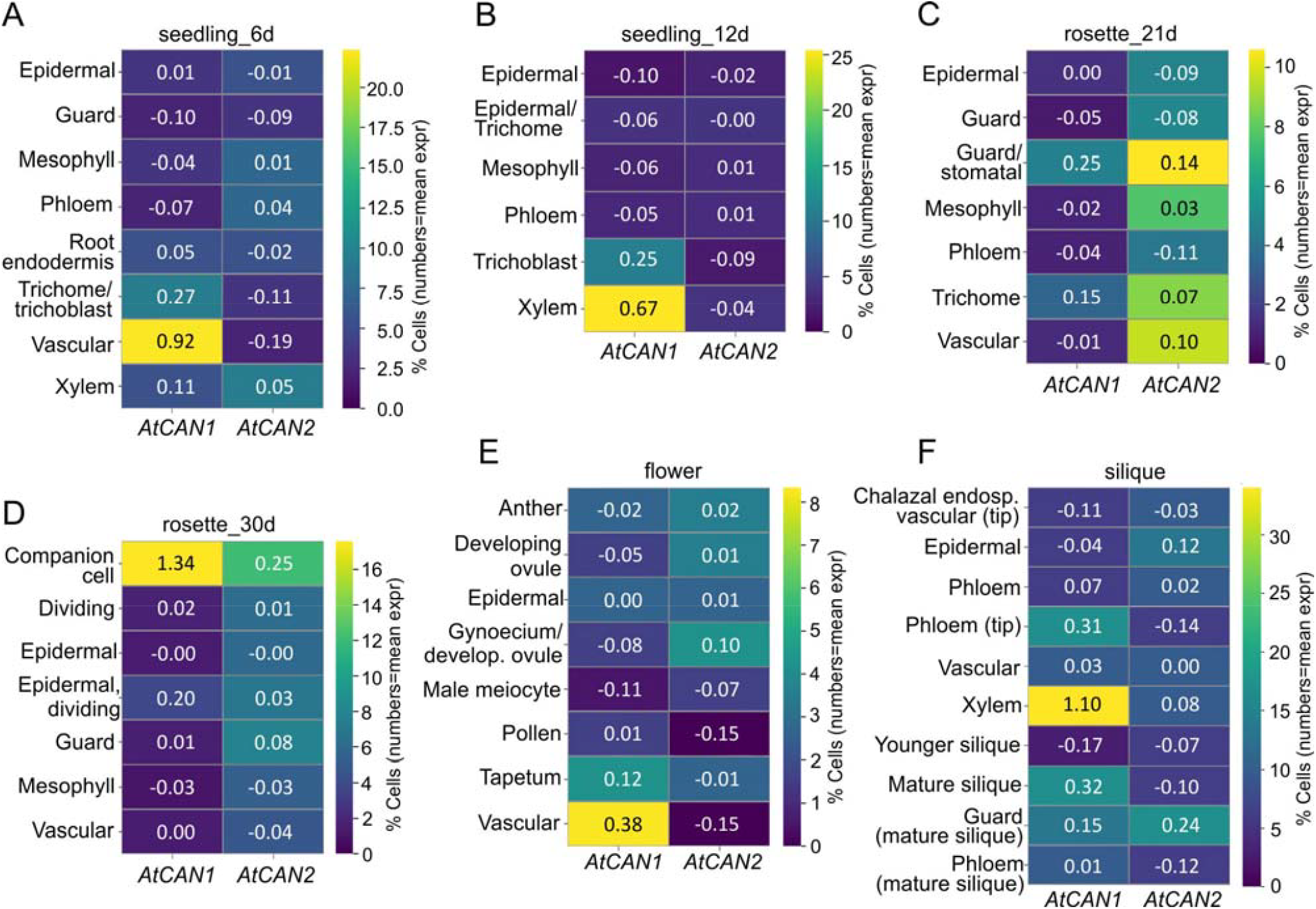
Expression of *AtCAN1* (AT3G56170) and *AtCAN2* (AT2G40410) based on single-nucleus RNA-seq data from the Arabidopsis Developmental Atlas [17]. Heat maps show the percentage of nuclei with detectable transcript levels across annotated cell types at six developmental stages: **(A)** Seedling 6 days, **(B)** Seedling 12 days, **(C)** Rosette 21 days, **(D)** Rosette 30 days, **(E)** Flower, and **(F)** Silique. Color intensity (viridis scale, 0% to stage-specific maximum) represents the percentage of nuclei per cell type in which expression was detected. Numbers within each cell indicate the mean normalized expression value for the respective gene and cell type. Color scales are independently normalized per panel (dynamic range and scale on the right). Cell type annotations follow the classification of the original atlas.

At the subsequent developmental stage, represented in the *Arabidopsis* Developmental Atlas by 21-day-old rosette leaves, elevated AtCAN1 activity remains detectable in trichomes and is also observed in cell populations classified as guard and stomatal lineages (**Figure 9C**). A partial increase in AtCAN2 gene expression was likewise detected in these cell types. In fully mature 30-day-old rosette leaves, a decrease in SNc gene activity is evident in guard cells, which is likely attributable to the completion of cell differentiation and the consequent cessation of protein accumulation encoded by both genes. In contrast, at this developmental stage, companion cells, which are a component of the vascular tissue, exhibited relatively high AtCAN1 expression and slightly lower AtCAN2 expression (**Figure 9D**).

In another plant organ, flowers, AtCAN1 expression was detected in cells annotated as vascular and, similarly to our GUS analyses, in the tapetum (**Figure 9E**). Furthermore, in siliques, a pronounced increase in AtCAN1 gene expression was observed during organ senescence, and its highest expression levels were recorded in cells defined as xylem (**Figure 9F**). It should be noted, however, that marker genes used in this database for xylem cell annotation, such as *IRX3/CESA7*, also play an essential role in the lignification of the endocarp of the silique valve [43, 44]. Therefore, cells of this tissue are likely also annotated as xylem cells.

In summary, data from the Arabidopsis Developmental Atlas largely confirm the key observations from our GUS reporter assays, which indicated *AtCAN1* expression in trichomes, guard cells, tapetum, vascular elements, and senescing siliques. Importantly, a novel finding of particular relevance to the discussion of AtCAN1 nuclease function is that, within vascular tissues, this protein is not only involved in xylem maturation but is also associated with the functioning of companion cells. The transcriptomic data analyzed here further support our general observation that, in most of the examined organs, expression of the gene encoding AtCAN2 is considerably less tissue-specific than that of AtCAN1.

### Phenotypic analysis of atcan1 insertion mutant lines

Among members of the SNc gene family, only the phenotype of Arabidopsis thaliana plants carrying a mutation in the AtCAN2 nuclease gene has been described to date. These plants were reported to exhibit enhanced tolerance to salt stress and reduced H_2_O_2_ accumulation [45]. Therefore, in the present study, we investigated the effects of AtCAN1 loss of function on plant phenotypic traits. To this end, seeds of a T-DNA insertion line carrying a mutation within the small fifth intron of the AtCAN1 gene were obtained from the SALK collection. Homozygous plants derived from this line were compared with wild-type *Arabidopsis thaliana* ecotype Columbia-0 (Col-0, WT). The most prominent phenotypic alteration observed in the *atcan1* mutant was a marked difference in rosette development. To validate this effect, the total rosette leaf area was quantified in 30-day-old plants. As shown in **Figure 10**, a statistically significant difference (p < 0.001) in rosette size was detected between the *atcan1* mutant and wild-type plants.

**Figure 10.**
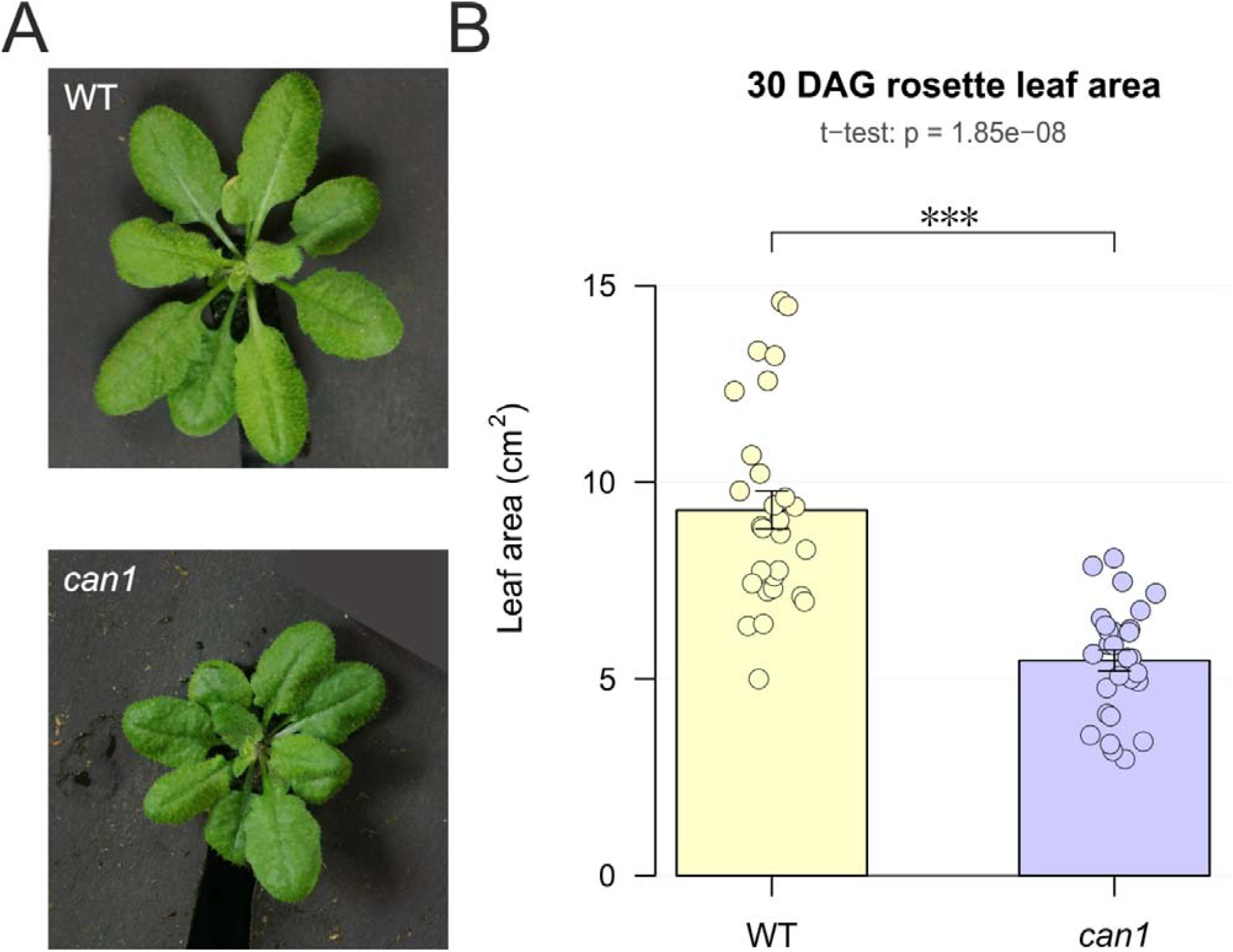
Rosette phenotype of wild-type and *AtCAN1* mutant plants. (A) Representative rosette images of wild-type (WT, top) and *can1* mutant (bottom) plants at 30 days after germination (DAG). (B) Quantification of total rosette leaf area. Bars represent mean ± SEM, individual data points show biological replicates (n = 28 per genotype). Statistical significance was determined by Welch’s t-test (***p < 0.001).

## Discussion

Plant development is closely associated with programmed cell death (PCD), with DNA degradation representing one of its key hallmarks. To date, research in this area has focused primarily on the role of nucleases belonging to the S1/P1 family. However, given the complexity of PCD at both the molecular and histological levels, multiple nucleases likely participate in these processes. During a survey of the *Arabidopsis thaliana* genome for genes encoding degradative nucleases, we found that, in addition to members of the S1/P1 family, *A. thaliana* also encodes two nucleases belonging to the SNc family. Previous studies have shown that, in bacteria, members of this family participate in various degradative processes [16,17]. Our earlier studies on plant representatives of this family confirmed the degradative nature of these enzymes, and analysis of publicly available microarray datasets suggested that they may be involved in specific types of plant PCD [12]. The expression profile of the *CrCAN* gene from *Citrus reticulata* also supports the hypothesis that this gene family is involved in PCD [13, 14].

To test this hypothesis, we performed histochemical analyses, considering GUS reporter assays to be particularly informative in this context. To minimize the possibility of technical artifacts that may occasionally accompany the GUS technique, we compared our observations with results from numerous transcriptomic studies using publicly available RNA-seq and microarray datasets. This analysis confirmed the validity of the observations obtained from our GUS-based experiments.

The results presented here demonstrate that the expression of *AtCAN1*, and to a lesser extent *AtCAN2*, is highly tissue-specific. The observed cases of their induction can be grouped into three main categories.

The first of these categories includes plant structures that are clear examples of organs undergoing PCD, such as the root cap, vascular bundle elements, senescent leaves, tapetum, and maturing seed pod. Among vegetative plant organs, the root cap is a classic example of a structure undergoing PCD. As the root grows and pushes through the soil, the outermost layer of the root cap, the lateral root cap cells (LRC), is subject to mechanical damage. Therefore, to protect the delicate apical meristem, these cells regularly undergo programmed cell death [46]. Because the GUS signal we observed closely correlates with the LRC, we conclude that the AtCAN1 nuclease is involved in PCD in this tissue. The correlation of GUS activity with the vascular bundles of the root, stem, and leaves also suggests a link between AtCAN1 and PCD, as xylogenesis in these cells represents another well-established example of programmed cell death.

Our research also demonstrated that the GUS signal correlates with the best characterized example of PCD in flowers, namely, tapetal cell death. During PCD, the tapetum releases enzymes, lipids, and nutrients that contribute to pollen grain formation. Our observations further revealed that the GUS marker driven by the *AtCAN1* promoter also follows this pattern. At later stages, as flower development progresses into seed formation, the GUS signal appeared two additional times. The first occurrence was in the transmitting tissue of the pistil style. Programmed cell death occurs in the transmitting tract around the time of pollination and facilitates the penetration of the pollen tube into the ovarian chamber. Moreover, it is believed that components of this tissue provide nutritional support to the pollen tube [47]. The final stage associated with seed formation, during which *AtCAN1* is expressed, occurred in the silique coats. As in the previous cases, the presence of PCD in the siliques was experimentally confirmed. This process is linked to silique valve dehiscence, which enables seed release. Silique cell death is commonly classified as a form of PCD-related senescence. It is accompanied by expression of the SAG13 gene, which is widely used as a marker for diverse processes associated with PCD [48].

The final and most striking evidence for the involvement of AtCAN1 nuclease in PCD is the detection of a GUS signal that emerges and spreads as leaf senescence progresses. Leaf senescence is one of the best-characterised forms of programmed cell death. During this process, gene expression shifts from pathways promoting photosynthesis and growth to those associated with degradation and recycling. This enables the plant to recover valuable nutrients (e.g., nitrogen and phosphorus from nucleic acids) and redistribute them to newly developing tissues or reproductive organs [49]. The involvement of the AtCAN1 nuclease in the redistribution of DNA degradation products is further supported by the high expression level of its gene in companion cells at the late rosette stage, as these cells are specialised in the active transport of metabolites into sieve tubes.

The second category of cases in which we identified *AtCAN1* promoter activity involves cells whose primary biological function is plant communication with the external environment. This group includes root hairs, stomatal guard cells, and hydathodes. Because these cells are particularly vulnerable to pathogens under natural conditions, they have evolved various protective mechanisms. Studies have shown that one of the main mechanisms employed by these structures is hypersensitive response (HR)-associated cell death [22–27]. In cells capable of HR, proteins involved in pathogen defence are pre-synthesized and maintained in a state ready for rapid activation, ensuring an immediate response to pathogen attack. Confirmation of the association of SNc genes with the response to biological stress is also provided by transcriptomic analyses, which show that the expression levels of these genes increase in other tissues following bacterial, fungal, and viral infections. Since, in known cases of HR, a rise in nucleolytic activity is observed, accompanied by host DNA degradation [28], we assume that the AtCAN1 gene is involved in this process. Moreover, it cannot be excluded that the activity of SNc nucleases in infection-exposed cells may target pathogen-derived genetic material, as has been demonstrated for PR-10 ribonuclease [56, 57].

A significant and unexpected finding of our study was the intense and highly specific expression of the *AtCAN1* promoter in a third group of plant structures, including stipules, trichomes, and the basal parts of the hypocotyl in early seedlings, as well as *AtCAN2* promoter activity in stipules. According to current knowledge, these structures, unlike the previously mentioned ones, are not subject to developmentally or environmentally induced programmed cell death in plants. Therefore, the presence of SNc nucleases in these structures cannot be explained by these processes. Instead, their expression might be associated with a shared feature of these structures – the occurrence of endoreduplication in their cells [58–60]. The involvement of the *AtCAN1* gene in endoreduplication is also supported by our transcriptomic data, which show a clear upward trend in its expression during induced endoreduplication.

Endoreduplication is a process that has long been observed in various tissues of higher plants. It results in endopolyploidy through repeated rounds of DNA synthesis uncoupled from cell division and is restricted to specific, terminally differentiated cells. The primary biological consequences of endoreduplication are cell enlargement and enhanced metabolic capacity, as the presence of multiple gene copies enables increased transcriptional activity. This functional interpretation is consistent with the frequent occurrence of endoreduplication in storage tissues, such as the endosperm of monocotyledons and the cotyledons of dicotyledonous plants, as well as in various secretory cells, including glandular trichomes. Although *Arabidopsis* trichomes are not classified as glandular, they also produce and accumulate specific secondary metabolites, mainly flavonoids, phenolics, and stress-related proteins [61]. By contrast, the presence of polyploidy in seedling stipules remains less readily explained. In *A*.*thaliana*, stipules are small, leaf-like structures located at the base of the leaf petiole and, unlike in many other species, do not play significant protective or mechanical roles. It has been suggested that polyploidy in these organs results from an endoreduplication-dependent mechanism that constrains tissue size [62].

However, regardless of the biological role of polyploidy, the question arises: what ultimately happens to the DNA accumulated in polyploid cells? Nucleic acids contain considerable amounts of nitrogen and phosphorus, which are well known to be limiting factors for plant growth due to their restricted availability in the environment. It is therefore widely accepted that the nucleolytic activity accompanying PCD in plants recycles these essential elements from dying cells. This raises the question of whether mechanisms exist to recover DNA components from polyploid cells once they have fulfilled their role as carriers of genetic information. Such a mechanism is likely to operate in the endosperm of monocot seeds, where strong nucleolytic activity has been observed during the mobilization of stored reserves [5, 63]. Nevertheless, no evidence of DNA recycling from living polyploid cells has been reported to date. In this context, our observation of SNc nuclease promoter activity in *Arabidopsis* polyploid cells is consistent with the hypothesis that plants may perform this type of recycling. Notably, because stipules, trichomes, and lower parts of the hypocotyl are not classical storage organs such as endosperm or cotyledons, the presumed degradation of polyploid DNA in these structures may represent a hitherto unidentified form of nutrient recycling. At this point, it is conceivable that polyploidy serves as a mechanism for storing components required for nucleic acid synthesis in growing leaves. Such a role could help to explain the activation of SNc nucleases observed in structures such as seedling stipules at the base of developing leaves.

The restricted expression pattern of AtCAN2, detected by GUS staining primarily in stipules and hydathodes, with only modest and often non-significant changes in the bulk transcriptomic datasets, is strikingly different from the broad and robust expression of AtCAN1. Both genes share substantial sequence similarity, but their divergent expression profiles suggest that the two paralogs have undergone subfunctionalization. The single-cell RNA-seq data reinforce this interpretation: whereas AtCAN1 shows clear enrichment in vascular and companion cells across multiple developmental stages, AtCAN2 expression is low and diffusely distributed, without strong cell-type specificity. Expression pattern indicates that AtCAN2 retains a residual role in PCD-related DNA degradation in a narrow set of contexts. Notably, the tissues in which AtCAN2 expression was detected — stipules and hydathodes — are sites where it is co-expressed with AtCAN1, suggesting that both paralogs may participate in the same processes: protection against pathogens in hydathodes and utilization of polyploid DNA in stipules. This co-expression pattern, together with the demonstrated involvement of the homologous CrCAN nuclease in PCD-related DNA degradation in secretory cavities of *Citrus grandis* ‘Tomentosa’ [13, 14, 64], supports a conserved role for SNc family members in tissue-specific nucleic acid turnover. However, beyond these shared expression domains, AtCAN2 appears to have diverged from AtCAN1 in its responsiveness to environmental cues, Sui et al. [45]: demonstrated that AtCAN2, but not AtCAN1, is transcriptionally responsive to salt stress. Conversely, in the pathogen infection datasets analyzed here, AtCAN1 showed significant upregulation upon challenge with *V. vulnificus, A. brassicicola*, TCV, and *F. oxysporum*, whereas AtCAN2 responses were generally weaker and less consistent (**Figure 7**). Taken together, these observations suggest that AtCAN2 may have been co-opted primarily for abiotic rather than biotic stress responses.

The relationship presented in this paper between SNc nuclease expression and programmed cell death indicates that at least two nuclease families, SNc and S1/P1, participate in genomic DNA degradation during this process. It is instructive to compare the tissue expression profiles of the SNc nucleases with those of the BFN1 protein, the best-studied representative of the S1/P1 family [10]. This comparison shows that nucleases of both families are expressed in senescent leaves, differentiating xylem, degrading tapetum, siliques, valves, and the transmitting tract. Interestingly, *AtCAN1* is also expressed in the root cap, a classic example of a tissue undergoing PCD, in which BFN1 expression has not been reported despite comprehensive studies of its promoter. Conversely, BFN1 expression was detected in several tissues in which AtCAN1 expression was not observed, such as the abscission zone, developing seed, and senescing petals. These analyses indicate that nuclease expression from both gene families overlaps in some classical PCD tissues, while retaining individual specificity in others. Notably, BFN1 expression was not detected in cells we considered potentially susceptible to HR-PCD due to their high exposure to pathogens, such as root hairs, stomata, and hydathodes. Moreover, a specific feature of AtCAN1 and AtCAN2 is their occurrence in polyploid cells, as *BFN1* expression has likewise not been observed in these elements to date.

Comparison of the expression patterns of SNc family proteins with those of BFN1 raises the question of whether any functional interdependence exists between these nucleases. Analysis of the mechanisms by which they operate within the cell may help address this issue. In young leaves, BFN1 was localized to filamentous reticulum-derived structures surrounding the nuclei, whereas in senescing cells it was detected within fragmented nuclei. This localization pattern supports the proposed role of BFN1 as a genomic DNA-degrading enzyme involved in senescence [10]. By contrast, our previous studies have shown that the nucleases described in this work, AtCAN1 and AtCAN2, are localized to the plasma membrane and exhibit partial homology to certain periplasmic components of bacterial ABC-type transporters. It is worth noting that the membrane localization of nucleases involved in PCD in plants has not been reported previously. However, bacterial homologues of these proteins are also anchored in the bacterial cell membrane [65]. Taking these observations into account, it can be concluded that S1/P1 nucleases are responsible for nuclear DNA degradation, whereas SNc nucleases couple the DNA degradation process with the extracellular transport of its breakdown products. Thus, nucleases from these two families appear to represent distinct roles in the cell. Their coexpression can therefore be interpreted as a form of cooperation, in which both types of activities complement each other. However, the question of whether SNc nucleases can function independently of S1/P1 nucleases remains unresolved, since among the known S1/P1 nucleases, only the expression profile of *BFN1* has been characterized. At the same time, this family in *A. thaliana* comprises four additional genes [11] whose biological functions remain to be determined.

The results presented suggest that the general function of SNc nucleases is to enhance the efficiency of utilization of DNA degradation products from different tissues at various stages of plant development. This hypothesis is consistent with the analysis of AtCAN1 mutant phenotypes, which showed that plants lacking the AtCAN1 protein exhibit reduced growth parameters, possibly due to impaired redistribution of nucleic acid building blocks. However, alternative explanations cannot be excluded. The strong vascular expression of AtCAN1 raises the possibility that the growth phenotype reflects compromised vascular function — including reduced nutrient or hormone transport — rather than a direct effect on nucleotide salvage. Additionally, xylem differentiation involves programmed cell death, and impaired clearance of nuclear debris in developing xylem elements could indirectly affect vascular conductivity. Determining the precise mechanism linking AtCAN1 loss to reduced leaf growth will require detailed cytological analysis of *atcan1* leaves and other organs, including measurements of cell number and cell size, as well as metabolomic profiling of nucleotide and nucleoside pools. In particular, given the correlation between AtCAN1 expression and endoreduplication observed in this study, it would be important to investigate whether endoreduplication-dependent processes in leaf cells contribute to the observed reduction in growth. Generation of *atcan1/atcan2* double mutants and crosses with S1/P1-type nuclease mutants would further clarify the extent of functional redundancy among DNA degradation pathways that contribute to plant growth.

## Conclusions

The results presented here demonstrate that the expression of AtCAN1 and AtCAN2 genes is highly tissue-specific. All observed cases of expression, revealed by promoter-driven GUS activity and transcriptomic analysis, fall into three main categories: (i) plant structures that clearly represent organs undergoing PCD, (ii) elements whose primary biological function is communication with the external environment and are therefore susceptible to hypersensitive response-associated cell death, and (iii) in polyploid cells. These results show that the involvement of SNc family nucleases in DNA degradation during various forms of plant PCD is as widespread as the previously reported involvement of S1/P1 proteins. In many cases, the expression profiles of nucleases from these two families overlap, suggesting cooperative action during PCD. Since SNc nucleases are plasma membrane proteins, whereas S1/P1 nucleases have been identified within nuclei, they appear to represent distinct yet complementary mechanisms of nucleolytic activity in the cell. Furthermore, we demonstrate that SNc nucleases are also specifically expressed in organs that do not undergo PCD but are characterized by endoreduplication. This suggests that SNc nucleases participate in a previously unexplored mechanism enabling the redistribution of polyploid DNA building blocks.

## We acknowledge

We thank Travis A. Lee and colleagues for providing the finalized annotated single-nucleus RNA-seq datasets from the Arabidopsis Developmental Atlas upon request. We thank Kotaro Torii and colleagues for providing single-cell RNA-seq data upon request.

## References

1. Van Durme M, Nowack MK. Mechanisms of developmentally controlled cell death in plants. Curr Opin Plant Biol. 2016;29:29–37. 10.1016/j.pbi.2015.10.013.

2. Balint-Kurti P. The plant hypersensitive response: concepts, control and consequences. Mol Plant Pathol. 2019;20:1163–78. 10.1111/mpp.12821.

3. Newton K, Dixit VM, Kayagaki N. Dying cells fan the flames of inflammation. Science (1979). 2021;374:1076–80. 10.1126/science.abi5934.

4. Sakamoto W, Takami T. Nucleases in higher plants and their possible involvement in DNA degradation during leaf senescence. J Exp Bot. 2014;65:3835–43. 10.1093/jxb/eru091.

5. Rewers M, Sliwinska E. Endoreduplication in the germinating embryo and young seedling is related to the type of seedling establishment but is not coupled with superoxide radical accumulation. J Exp Bot. 2014;65:4385–96. 10.1093/jxb/eru210.

6. Lesniewicz K, Pieńkowska J, Poreba E. Characterization of nucleases involved in seedling development of cauliflower. J Plant Physiol. 2010;167:1093–100. 10.1016/j.jplph.2010.03.011.

7. Lambert R, Quiles FA, Cabello-Díaz JM, Piedras P. Purification and identification of a nuclease activity in embryo axes from French bean. Plant Sci. 2014;224:137–43. 10.1016/j.plantsci.2014.04.017.

8. Delgado-García E, Piedras P, Gómez-Baena G, García-Magdaleno IM, Pineda M, Gálvez-Valdivieso G. Nucleoside Metabolism Is Induced in Common Bean During Early Seedling Development. Front Plant Sci. 2021;12. 10.3389/fpls.2021.651015.

9. Perez-Amador MA, Abler ML, De Rocher EJ, Thompson DM, Van Hoof A, LeBrasseur ND, et al. Identification of BFN1, a bifunctional nuclease induced during leaf and stem senescence in Arabidopsis. Plant Physiol. 2000;122:169–79. 10.1104/pp.122.1.169.

10. Farage-Barhom S, Burd S, Sonego L, Perl-Treves R, Lers A. Expression analysis of the BFN1 nuclease gene promoter during senescence, abscission, and programmed cell death-related processes. J Exp Bot. 2008;59:3247– 58. 10.1093/jxb/ern176.

11. Lesniewicz K, Karlowski WM, Pienkowska JR, Krzywkowski P, Poreba E. The plant s1-like nuclease family has evolved a highly diverse range of catalytic capabilities. Plant Cell Physiol. 2013;54:1064–78. 10.1093/pcp/pct061.

12. Lesniewicz K, Poreba E, Smolarkiewicz M, Wolff N, Stanisławski S, Wojtaszek P. Plant plasma membranebound staphylococcal-like DNases as a novel class of eukaryotic nucleases. BMC Plant Biol. 2012;12:195. 10.1186/1471-2229-12-195.

13. Liang M, Huai B, Lin J, Liang X, He H, Bai M, et al. Ca2+-and Zn2+-dependent nucleases co-participate in nuclear DNA degradation during programmed cell death in secretory cavity development in Citrus fruits. Tree Physiol. 2024;44. 10.1093/treephys/tpad122.

14. Liang M, Huai B, Lin J, He H, Bai M, Wu H. NAC transcription factors regulate Ca2+-dependent and Zn2+ dependent nucleases to cooperatively participate in nuclear DNA degradation during programmed cell death in secretory cavity cells of Citrus fruits. Sci Hortic. 2026;357. 10.1016/j.scienta.2026.114642.

15. Weigel Detlef, Glazebrook Jane. Arabidopsisl?: a laboratory manual. Cold Spring Harbor Laboratory Press; 2002.

16. Yu Y, Zhang H, Long Y, Shu Y, Zhai J. Plant Public RNA-seq Database: a comprehensive online database for expression analysis of ~45 000 plant public RNA-Seq libraries. Plant Biotechnol J. 2022;20:806–8. 10.1111/pbi.13798.

17. Lee TA, Illouz-Eliaz N, Nobori T, Xu J, Jow B, Nery JR, et al. A single-cell, spatial transcriptomic atlas of the Arabidopsis life cycle. Nat Plants. 2025;11:1960–75. 10.1038/s41477-025-02072-z.

18. Wolf FA, Angerer P, Theis FJ. SCANPY: Large-scale single-cell gene expression data analysis. Genome Biol. 2018;19. 10.1186/s13059-017-1382-0.

19. Easlon HM, Bloom AJ. Easy Leaf Area: Automated digital image analysis for rapid and accurate measurement of leaf area. Appl Plant Sci. 2014;2. 10.3732/apps.1400033.

20. Birnbaum K, Shasha DE, Wang JY, Jung JW, Lambert GM, Galbraith DW, et al. A Gene Expression Map of the Arabidopsis Root. Science (1979). 2003;302:1956–60. 10.1126/science.1090022.

21. Karve R, Suárez-Román F, Iyer-Pascuzzi AS. The transcription factor NIN-LIKE PROTEIN7 controls border-like cell release. Plant Physiol. 2016;171:2101–11. 10.1104/pp.16.00453.

22. Becker JD, Takeda S, Borges F, Dolan L, Feijó JA. Transcriptional profiling of Arabidopsis root hairs and pollen defines an apical cell growth signature. BMC Plant Biol. 2014;14:197. 10.1186/s12870-014-0197-3.

23. Klepikova A V., Kasianov AS, Gerasimov ES, Logacheva MD, Penin AA. A high resolution map of the Arabidopsis thaliana developmental transcriptome based on RNA-seq profiling. Plant Journal. 2016;88:1058–70. 10.1111/tpj.13312.

24. Kondo Y, Fujita T, Sugiyama M, Fukuda H. A novel system for xylem cell differentiation in arabidopsis thaliana. Mol Plant. 2015;8:612–21. 10.1016/j.molp.2014.10.008.

25. Wendrich JR, Möller BK, Li S, Saiga S, Sozzani R, Benfey PN, et al. Framework for gradual progression of cell ontogeny in the Arabidopsis root meristem. Proc Natl Acad Sci U S A. 2017;114:E8922–9. 10.1073/pnas.1707400114.

26. Robson JK, Tidy AC, Thomas SG, Wilson ZA. Environmental regulation of male fertility is mediated through Arabidopsis transcription factors bHLH89, 91, and 10. J Exp Bot. 2024;75:1934–47. 10.1093/jxb/erad480.

27. Zhu E, You C, Wang S, Cui J, Niu B, Wang Y, et al. The DYT1-interacting proteins bHLH010, bHLH089 and bHLH091 are redundantly required for Arabidopsis anther development and transcriptome. Plant Journal. 2015;83:976–90. 10.1111/tpj.12942.

28. Li DD, Xue JS, Zhu J, Yang ZN. Gene regulatory network for tapetum development in arabidopsis thaliana. Front Plant Sci. 2017;8. 10.3389/fpls.2017.01559.

29. Adrian J, Chang J, Ballenger CE, Bargmann BOR, Alassimone J, Davies KA, et al. Transcriptome dynamics of the stomatal lineage: Birth, amplification, and termination of a self-renewing population. Dev Cell. 2015;33:107–18. 10.1016/j.devcel.2015.01.025.

30. Lee LR, Wengier DL, Bergmann DC. Cell-type–specific transcriptome and histone modification dynamics during cellular reprogramming in the Arabidopsis stomatal lineage. Proc Natl Acad Sci U S A. 2019;116:21914–24. 10.1073/pnas.1911400116.

31. Schliep M, Ebert B, Simon-Rosin U, Zoeller D, Fisahn J. Quantitative expression analysis of selected transcription factors in pavement, basal and trichome cells of mature leaves from Arabidopsis thaliana. Protoplasma. 2010;241:29–36. 10.1007/s00709-009-0099-7.

32. Park YS, Kim SK, Kim SY, Kim KM, Ryu CM. The transcriptome analysis of the Arabidopsis thaliana in response to the Vibrio vulnificus by RNA-sequencing. PLoS One. 2019;14. 10.1371/journal.pone.0225976.

33. Fallath T, Kidd BN, Stiller J, Davoine C, Björklund S, Manners JM, et al. MEDIATOR18 and MEDIATOR20 confer susceptibility to Fusarium oxysporum in Arabidopsis thaliana. PLoS One. 2017;12. 10.1371/journal.pone.0176022.

34. Wu C, Li X, Guo S, Wong SM. Analyses of RNA-Seq and sRNA-Seq data reveal a complex network of anti-viral defense in TCV-infected Arabidopsis thaliana. Sci Rep. 2016;6. 10.1038/srep36007.

35. Torii K, Inoue K, Bekki K, Haraguchi K, Kubo M, Kondo Y, et al. A guiding role of the Arabidopsis circadian clock in cell differentiation revealed by time-series single-cell RNA sequencing. Cell Rep. 2022;40. 10.1016/j.celrep.2022.111059.

36. Ding A, Xu C, Xie Q, Zhang M, Yan N, Dai C, et al. ERF4 interacts with and antagonizes TCP15 in regulating endoreduplication and cell growth in Arabidopsis. J Integr Plant Biol. 2022;64:1673–89. 10.1111/jipb.13323.

37. de Luxán-Hernández C, Lohmann J, Tranque E, Chumova J, Binarova P, Salinas J, et al. MDF is a conserved splicing factor and modulates cell division and stress response in Arabidopsis. Life Sci Alliance. 2023;6. 10.26508/lsa.202201507.

38. Schwarz EM, Roeder AHK. Transcriptomic effects of the cell cycle regulator LGO in Arabidopsis sepals. Front Plant Sci. 2016;7. 10.3389/fpls.2016.01744.

39. López-Juez E, Dillon E, Magyar Z, Khan S, Hazeldine S, De Jager SM, et al. Distinct light-initiated gene expression and cell cycle programs in the shoot apex and cotyledons of Arabidopsis. Plant Cell. 2008;20:947–68. 10.1105/tpc.107.057075.

40. Yagi H, Tamura K, Matsushita T, Shimada T. Spatiotemporal relationship between auxin dynamics and hydathode development in Arabidopsis leaf teeth. Plant Signal Behav. 2021;16. 10.1080/15592324.2021.1989216.

41. Smyth DR, Bowman JL, Meyerowitz EM. Early flower development in Arabidopsis. Plant Cell. 1990;2:755–67. 10.2307/3869174.

42. Jakoby MJ, Falkenhan D, Mader MT, Brininstool G, Wischnitzki E, Platz N, et al. Transcriptional profiling of mature Arabidopsis trichomes reveals that NOECK encodes the MIXTA-like transcriptional regulator MYB106. Plant Physiol. 2008;148:1583–602. 10.1104/pp.108.126979.

43. Mitsuda N, Ohme-Takagi M. NAC transcription factors NST1 and NST3 regulate pod shattering in a partially redundant manner by promoting secondary wall formation after the establishment of tissue identity. Plant Journal. 2008;56:768–78. 10.1111/j.1365-313X.2008.03633.x.

44. Barros J, Serk H, Granlund I, Pesquet E. The cell biology of lignification in higher plants. Annals of Botany. 2015;115:1053–74. 10.1093/aob/mcv046.

45. Sui W, Guo K, Li L, Liu S, Takano T, Zhang X. Arabidopsis Ca2+-dependent nuclease AtCaN2 plays a negative role in plant responses to salt stress. Plant Science. 2019;281:213–22. 10.1016/j.plantsci.2018.12.007.

46. Kumar N, Iyer-Pascuzzi AS. Shedding the last layer: Mechanisms of root cap cell release. Plants. 2020;9. 10.3390/plants9030308.

47. Crawford BCW, Ditta G, Yanofsky MF. The NTT Gene Is Required for Transmitting-Tract Development in Carpels of Arabidopsis thaliana. Current Biology. 2007;17:1101–8. 10.1016/j.cub.2007.05.079.

48. Dhar N, Caruana J, Erdem I, Subbarao K V., Klosterman SJ, Raina R. The Arabidopsis senescencE-associated gene 13 regulates dark-induced senescence and plays contrasting roles in defense against bacterial and fungal pathogens. Molecular Plant-Microbe Interactions. 2020;33:754–66. 10.1094/MPMI-11-19-0329-R.

49. Guo Y, Ren G, Zhang K, Li Z, Miao Y, Guo H. Leaf senescence: progression, regulation, and application. Molecular Horticulture. 2021;1. 10.1186/s43897-021-00006-9.

50. Hogg B V., Kacprzyk J, Molony EM, O’Reilly C, Gallagher TF, Gallois P, et al. An in vivo root hair assay for determining rates of apoptotic-like programmed cell death in plants. Plant Methods. 2011;7. 10.1186/1746-4811-7-45.

51. Ye W, Munemasa S, Shinya T, Wu W, Ma T, Lu J, et al. Stomatal immunity against fungal invasion comprises not only chitin-induced stomatal closure but also chitosan-induced guard cell death. Proceedings of the National Academy of Sciences. 2020;117:20932–42. 10.1073/pnas.1922319117.

52. Yang LN, Liu H, Wang YP, Seematti J, Grenville-Briggs LJ, Wang Z, et al. Pathogen-Mediated Stomatal Opening: A Previously Overlooked Pathogenicity Strategy in the Oomycete Pathogen Phytophthora infestans. Front Plant Sci. 2021;12. 10.3389/fpls.2021.668797.

53. Taks NW, van Hulten M, van Splunter-Berg JA, Chatterjee S, Stevens FD, Paauw M, et al. Arabidopsis CNL receptor SUT1 confers immunity in hydathodes against the vascular pathogen Xanthomonas campestris pv. campestris. PLoS Pathog. 2025;21. 10.1371/journal.ppat.1013256.

54. Kobae Y, Sekino T, Yoshioka H, Nakagawa T, Martinoia E, Maeshima M. Loss of AtPDR8, a plasma membrane ABC transporter of Arabidopsis thaliana, causes hypersensitive cell death upon pathogen infection. Plant Cell Physiol. 2006;47:309–18. 10.1093/pcp/pcj001.

55. Chiusano ML, Incerti G, Colantuono C, Termolino P, Palomba E, Monticolo F, et al. Arabidopsis thaliana response to extracellular dna: Self versus nonself exposure. Plants. 2021;10. 10.3390/plants10081744.

56. Park CJ, Kim KJ, Shin R, Park JM, Shin YC, Paek KH. Pathogenesis-related protein 10 isolated from hot pepper functions as a ribonuclease in an antiviral pathway. Plant Journal. 2004;37:186–98. 10.1046/j.1365-313X.2003.01951.x.

57. Röck M, Heel SV, Juen FS, Eidelpes R, Kreutz C, Breuker K, et al. The PR-10 Protein Pru p 1 is an Endonuclease that Preferentially Cleaves Single–Stranded RNA. ChemBioChem. 2024;25. 10.1002/cbic.202400204.

58. Vuolo F, Kierzkowski D, Runions A, Hajheidari M, Mentink RA, Gupta M Das, et al. LMI1 homeodomain protein regulates organ proportions by spatial modulation of endoreduplication. Genes Dev. 2018;32:1361–6. 10.1101/gad.318212.118.

59. Walker JD, Oppenheimer DG, and JC, Larkin JC. SIAMESE, a gene controlling the endoreduplication cell cycle in Arabidopsis thaliana trichomes. Development. 2000;127:3931–40. 10.1242/dev.127.18.3931.

60. Tanaka R, Amijima M, Iwata Y, Koizumi N, Mishiba K ichiro. Effect of light and auxin transport inhibitors on endoreduplication in hypocotyl and cotyledon. Plant Cell Rep. 2016;35:2539–47. 10.1007/s00299-016-2054-3.

61. Huebbers JW, Mantz M, Panstruga R, Huesgen PF. Proteomics dataset on detached and purified Arabidopsis thaliana rosette leaf trichomes. Data Brief. 2023;46. 10.1016/j.dib.2023.108897.

62. Jacqmard A, De Veylder L, Segers G, Almeida Engler J, Bernier G, Van Montagu M, et al. Expression of CKS1At in Arabidopsis thaliana indicates a role for the protein in both the mitotic and the endoreduplication cycle. Planta. 1999;207:496–504. 10.1007/s004250050509.

63. Spanò C, Buselli R, Ruffini Castiglione M, Bottega S, Grilli I. RNases and nucleases in embryos and endosperms from naturally aged wheat seeds stored in different conditions. J Plant Physiol. 2007;164:487–95. 10.1016/j.jplph.2006.03.015.

64. Bai M, Liang M, Huai B, Gao H, Tong P, Shen R, et al. Ca2+-dependent nuclease is involved in DNA degradation during the formation of the secretory cavity by programmed cell death in fruit of Citrus grandis ‘Tomentosa.’ J Exp Bot. 2020; April. 10.1093/jxb/eraa199.

65. Kiedrowski MR, Crosby HA, Hernandez FJ, Malone CL, McNamara JO, Horswill AR. Staphylococcus aureus Nuc2 is a functional, surface-attached extracellular nuclease. PLoS One. 2014;9. 10.1371/journal.pone.0095574.

